# VIRAL DELIVERY OF GDNF PROMOTES FUNCTIONAL INTEGRATION OF HUMAN STEM CELL GRAFTS IN PARKINSON’S DISEASE

**DOI:** 10.1101/870725

**Authors:** Carlos W Gantner, Isabelle R de Luzy, Jessica A Kauhausen, Niamh Moriarty, Jonathan C Niclis, Vanessa Penna, Cameron P. Hunt, Christopher R Bye, Charlotte M Ermine, Colin W Pouton, Deniz Kirik, Lachlan H Thompson, Clare L Parish

**Author notes:** Author for correspondence: Clare Parish.

## Abstract

The derivation of neurotransmitter and region-specific neuronal populations from human pluripotent stem cells (PSC) provides impetus for advancing cell therapies into the clinic. At the forefront is our ability to generate ventral midbrain (VM) dopaminergic (DA) progenitors, suitable for transplantation in Parkinson’s disease (PD). Pre-clinical studies, however, have highlighted the low proportion of DA neurons within these grafts and their inferior plasticity by comparison to human fetal donor transplants. Here we sought to examine whether modification of the host environment, through viral delivery of a developmentally critical molecule, glial cell line-derived neurotrophic factor (GDNF), could improve graft survival, integration and function in Parkinsonian rodents. Utilising LMX1A- and PITX3-GFP hPSC reporter lines, we tracked the response of DA progenitors implanted into either a GDNF-rich environment, or in a second group, after a 3-week delay in onset of exposure. We found that early exposure of the graft to GDNF promoted survival of DA and non-DA cells, leading to enhanced motor recovery in PD rats. Delayed overexpression of intrastriatal GDNF also promoted motor recovery in transplanted rats, through alternate selective mechanisms including enhanced A9/A10 specification, increased DA graft plasticity, greater activation of striatal neurons and elevated DA metabolism. Lastly, transcriptional profiling of the grafts highlighted novel genes underpinning these changes. Collectively these results demonstrate the potential of targeted neurotrophic gene therapy strategies to improve human PSC graft outcomes.

## Introduction

Clinical trials have provided evidence that human ventral midbrain (VM) dopamine (DA) progenitors, transplanted into the denervated striatum of Parkinson’s disease (PD) patients, can structurally integrate, restore DA transmission and alleviate motor symptoms for decades [reviewed in ^1^]. Hindering this therapy has been the reliance on human fetal tissue that, in addition to availability and ethical constraints, remains poorly standardized. Consequently, focus has shifted to generating replacement DA progenitors from human pluripotent stem cells (hPSC). hPSC, derived either from the inner cell mass of the developing blastocyst (ESC – embryonic stem cells) or by reprogramming of somatic cells by defined factors (iPSC – induced pluripotent stem cells), are capable of generating derivatives of all three germ layers *in vitro*, including DA progenitors. Employing highly standardised procedures, that encompass modelling the developing VM environment and using both tissue-appropriate laminin substrates and extrinsic molecules, recent years have seen rapid advancements in protocols for the generation of *bona fide* VM DA neurons from both human embryonic and induced pluripotent stem cells ^2-4^.

Following transplantation, human PSC-derived DA progenitors are capable of maturing into DA neurons, adopting appropriate target acquisition and restoring gross motor deficits ^5, 6^. However, overall proportions of DA neurons within the grafts remain low. In part, this likely reflects the poor survival of these progenitors/neurons during and immediately following transplantation, as reported for fetal tissue grafts ^7, 8^. In addition, transplanted hESC-derived DA neurons show inferior axonal growth, compared to human fetal VM donor tissue. For example, Grealish et al., (2014), employing a human specific neural cell adhesion molecule and tyrosine hydroxylase (TH) co-immunoreactivity to selectively compare graft-derived DA axons from human ESC-derived and human fetal donor grafts, demonstrated reduced innervation of host A9-target tissue (dorsolateral and dorsomedial striatum) from the ESC-derived grafts. Such observations suggest that greater efforts to promote survival and axonal plasticity of human PSC-derived DA neurons following transplantation are required to improve outcomes after grafting and make this therapy clinically competitive.

Unlike the developing brain that contains a cocktail of neurotrophic and morphogenic proteins that influence survival, differentiation and axonal connectivity, the adult brain is a notably less permissive environment for the survival and integration of transplanted progenitors, with many developmental cues downregulated or absent. First identified for its role in the survival and plasticity of embryonic midbrain dopamine neurons in culture ^9^, glial cell line-derived neurotrophic factor (GDNF) has been shown to increase the survival, plasticity and metabolism of DA neurons in pre-clinical and clinical studies for Parkinson’s disease – see reviews ^10-12^. Recombinant GDNF protein has also been used to promote the survival of DA progenitors within fetal donor preparations prior to transplantation ^13^, with studies in rodents and non-human primates demonstrating the benefits of prolonged delivery into the host tissue (via infusion ^14-17^, or injection of viral vectors ^18-24^ for promoting survival, plasticity and functionally appropriate integration of DA-rich fetal VM tissue grafts.

Here we have assessed the impact of long-term GDNF overexpression on human PSC-derived VM DA progenitor grafts. We showed that exposure of the graft to GDNF had a significant impact on survival, plasticity and functional integration and notably that the timing of exposure was an important variable. Using a recombinant adeno-associated viral (AAV) vector to overexpress GDNF in the denervated host striatum, we showed that transplanting hESC-derived DA progenitors directly into a GDNF-rich environment promoted survival yet impacted on the capacity of the graft to innervate the host striatum. In contrast, exposure of the DA progenitors to GDNF several weeks after implantation had no effect on survival but positively impacted on DA neuron maturation, innervation of DA-relevant targets and functional outcomes. With human PSC therapies rapidly moving towards the clinic, these findings have important implications for promoting survival, plasticity, integration and thereby function of human PSC derived DA grafts, targeted at restoring DA transmission in PD patients.

## Results

### Validation of DA differentiation, GDNF expression and GDNF-induced motor recovery

Midbrain DA neurons were differentiated from a human embryonic stem cell line expressing enhanced green fluorescent protein under the PITX3 promoter (PITX3-GFP) according to a recent, clinically relevant xenogeneic-free protocol (Figure1A) ^2^. Enrichment of DA progenitors was confirmed by OTX2+/FOXA2+ co-expression at 15 days in vitro (DIV) (Figure 1B), accounting for >85% cells in culture, with limited numbers of PITX2, PAX6 or BARHL1+ cells observed (indicative of unintended lateral or rostral midbrain contaminants) (Figure 1C). While progenitors were isolated at 20 DIV for transplantation, parallel cultures, extended to 25DIV, confirmed the ability of the cells to generate *bona fide* VM dopamine neurons, displaying high proportions of GFP (PITX3), TH and FOXA2 co-expression, as previously described^2^, (Figure 1E). Quantitative PCR analysis of maturing DA progenitors *in vitro* confirmed upregulation of early and late DA determinant genes LMX1A and TH, respectively, and importantly the expression of the canonical GDNF receptor, GFRα1 (Figure 1D). Maintenance of Ret expression within VM progenitors (also expressed on human PSCs^3^), was confirmed by real-time PCR (Supplementary Figure 1), indicating that the VM progenitors were capable of responding to GDNF at the time of transplantation. Long term stable expression of AAV5-GDNF (Figures 1F and Supplementary Figure 2) and the control AAV5-mCherry (Supplementary Figure S3B) were observed at 6 months within the host striatum, as well as along the ipsilateral striatonigral tract, using GDNF and RFP immunohistochemistry, respectively.

**Figure 1:**
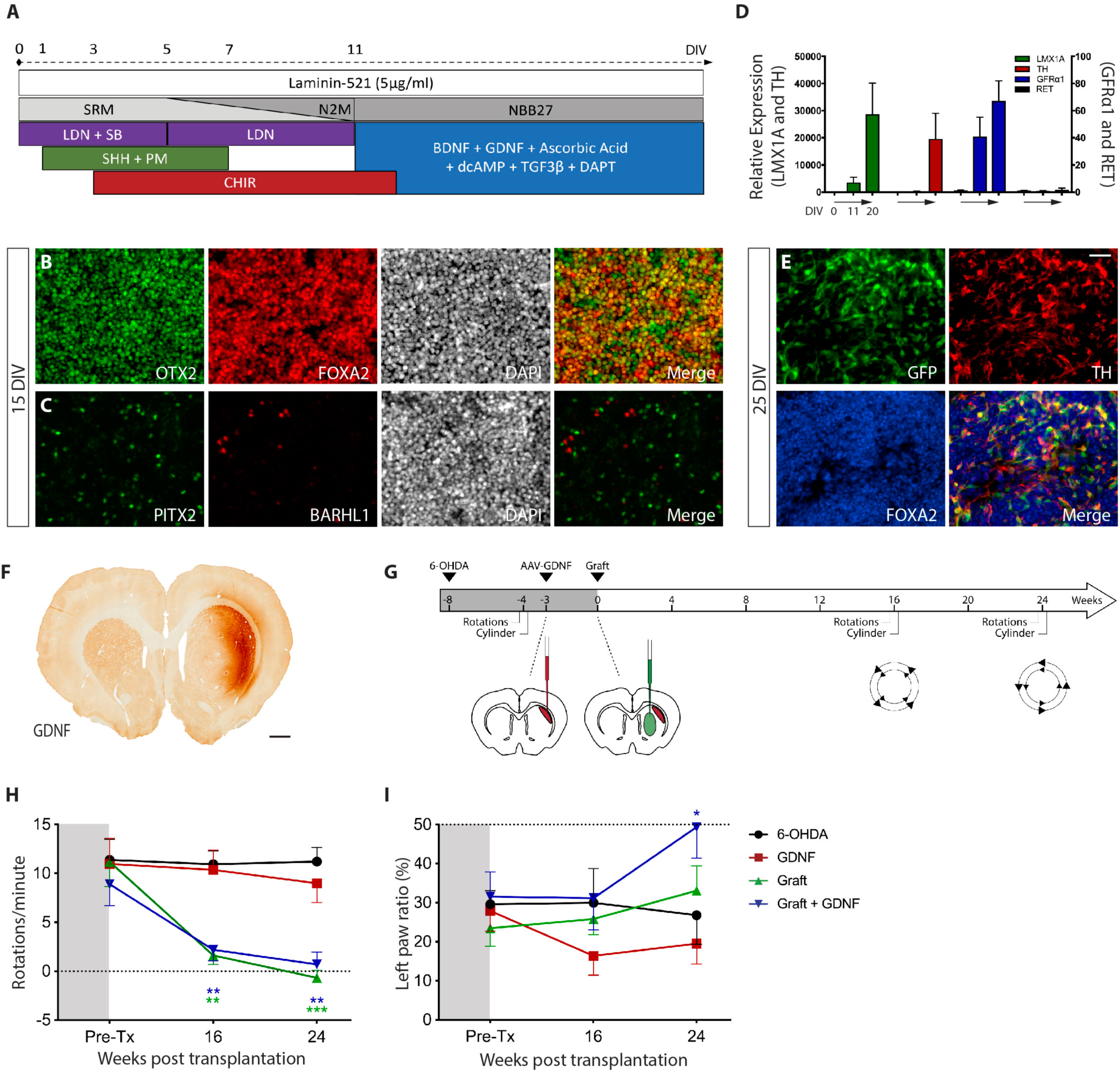
Exposure of human ESC-derived DA transplants to GDNF, from the time of implantation, improved motor deficits in Parkinsonian rats. (A) Differentiation protocol for human ESCs into VM progenitors, suitable for transplantation. (B) Photomicrographs showing high proportions of OTX2 and FOXA2 co-expression within progenitors at 15DIV. DIV, days in vitro. (C) Low numbers of PITX2+ and BARHL1+ cells, indicative of off-target rostral/forebrain progenitor populations, validated efficiency of the VM differentiation. (D) Graph showing mRNA levels of the pluripotency marker OCT4, VM progenitor/neuron markers (LMX1A, NURR1, TH) and GDNF receptors (Ret, GFRα1) during differentiation, relative to undifferentiated human ESCs. All values normalised to HPRT1 expression. n = 3 independent cultures. (E) Validation of VM differentiation into DA neurons, as confirmed by PITX3-GFP, TH and FOXA2 expression at 25DIV. (F) Coronal section of the rat brain illustrating sustained GDNF expression 6 months after intrastriatal injection of AAV-GDNF. (G) Schematic summary of *in vivo* study, inclusive of behavioural testing, GDNF viral delivery and cell transplantation. (H) All grafted animals showed complete reversal of amphetamine-induced rotations at 24 weeks (I) Correction of forelimb asymmetry in the cylinder test was only observed in rats receiving grafts in the presence of GDNF (blue line). (B,C,E) Scale bar, 50µm. (F) Scale bar, 1mm. Lesion, n = 11; GDNF, n = 7; Graft, n = 7 and Graft+GDNF, n = 7. Two-way ANOVA compared to 6-OHDA with Dunnett correction for multiple comparisons.

**Figure 2:**
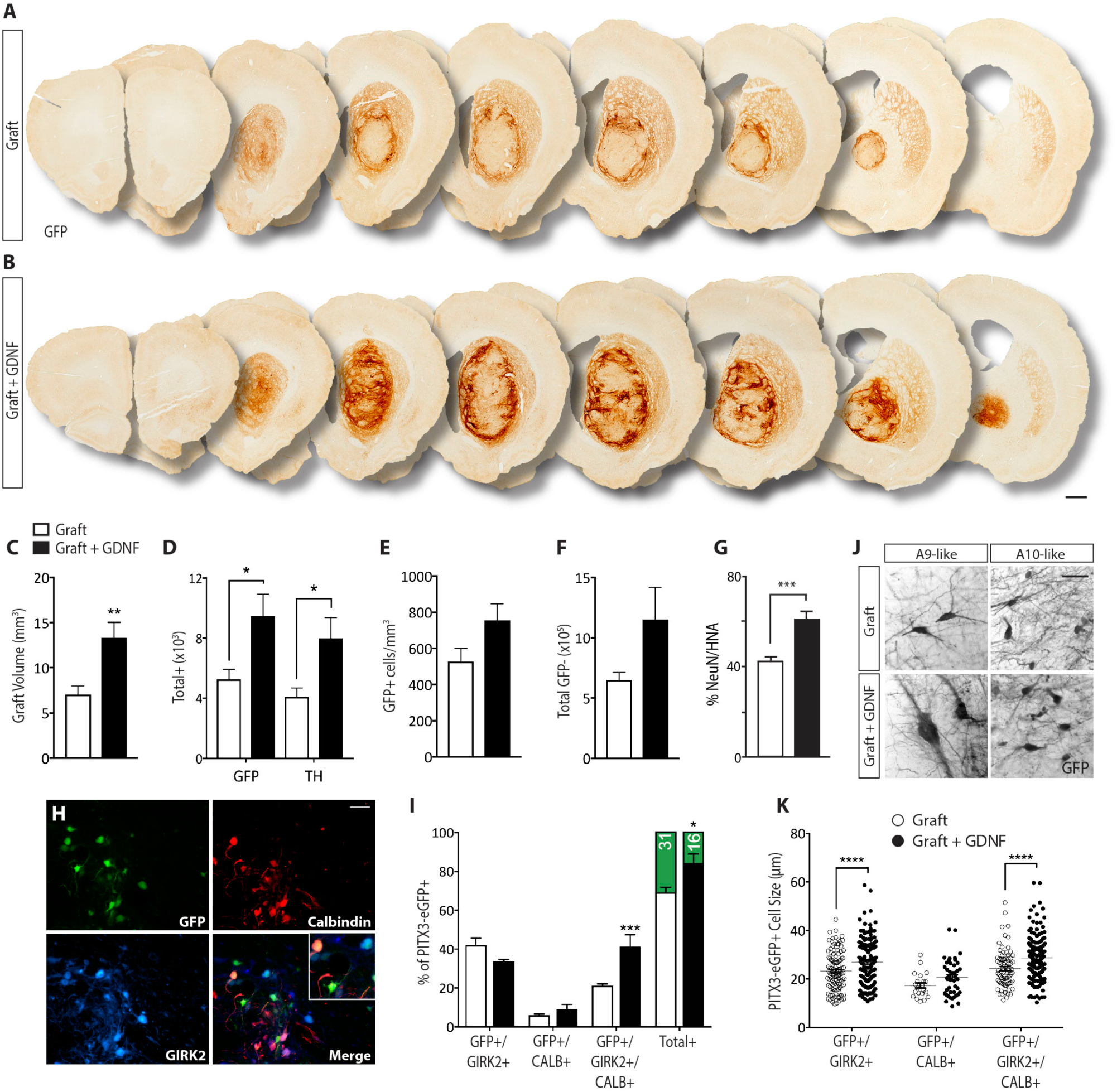
Implantation of human PSC-derived DA progenitors into a GDNF-rich environment promotes graft survival. (A-B) Representative overview of a hPSC-derived VM progenitor graft in the absence (A) and presence (B) of AAV-GDNF, showing PITX3-GFP expression. Scale bar, 1mm. (C) Assessment of graft volume showed significantly larger grafts in the presence of GDNF. (D-E) GFP+ and TH+ cell numbers commensurately increases with volume in animals receiving transplants in the presence of GDNF (D), resulting in unchanged GFP+ cell density (E). (F) Exposure of the graft to GDNF also increased the number of GFP-cells within the transplants. (G) Cell + GDNF grafts showed enhanced maturation, as revealed by the increased proportion of NeuN+ neurons within the graft. (H-I) Representative image showing GFP (PITX3) cells co-expressing GIRK2 and/or Calbindin (CALB), indicative of A9-like and A10-like VM DA neurons (G), and their relative contribution (H). Note, +/+ represents GFP+GIRK2+CALB+ co-immunoreactive cells. Green bar represents % of GFP+ cells that are neither GIRK2+ or CALB+. Scale bar, 50µm. (J) Representative images of A9-like (large, angled soma, commonly found at the graft periphery) or A10-like (small, circular soma, found predominantly in the graft core) ± GDNF. (K) GFP/GIRK co-expressing neurons were significantly larger in the presence of GDNF suggestive of increased maturation. Data in panels C-F,H,J expressed as mean ± SEM, Students t-test (n = 7/group).

Rats receiving 6-OHDA lesions alone (6-OHDA), Lesion + AAV-GDNF (GDNF), Lesion + PITX3-GFP VM DA progenitor transplants (Graft) or Lesion + combined AAV-GDNF and PITX3-GFP VM DA graft (Graft + GDNF) underwent a battery of motor tests prior to grafting and at 24 weeks post-transplantation (Figure 1G). Note, all rats included in the study showed an amphetamine induced rotational asymmetry of ≥5 rotations/min pre-transplantation, that was persistent and stable for 24 weeks in the 6-OHDA lesion control group (Figure1H, blackline). As expected, expression of GDNF from an AAV vector in the absence of a cell transplant did not reverse rotational behaviour in the chronically lesioned animals (Figure1H, red line). In contrast, rats receiving a Graft with or without GDNF showed complete recovery by 24 weeks (Figure 1H, green and blue lines, respectively). In the cylinder test (an assessment of spontaneous motor asymmetry), motor dysfunction was reflected as a decrease in weight-bearing contralateral paw touches following lesioning (Figures 1I, black line). Recovery was only observed in the Graft + GDNF group at 24 weeks (Figure 1I, blue line). The time period of recovery (6 months) observed here highlights the necessity of extended observational studies, as previously described for both human fetal ^25^ and human PSC-derived DA progenitor transplants ^5^.

### Transplantation of PSC-derived DA precursors into a GDNF-overexpressing environment promotes graft survival but disrupts axonal targeting

At 24 weeks after transplantation, all animals (± GDNF) displayed surviving grafts, as confirmed by GFP expression (Figures 2A and 2B). Importantly, we found no gross evidence of aberrant graft growth, i.e. no cell overgrowth or distortion of host tissue. This was supported by the extremely low proportion of Ki67+ cells within the grafts at 24 weeks, suggesting that the *in vitro* differentiation and subsequent maturation of the pluripotent stem cell-derived progenitors *in situ* was sufficient to deplete proliferative pools. In all animals the GFP+ grafts were predominantly confined to the striatum. The validity of PITX3-eGFP as an accurate marker of dopaminergic neurons was confirmed by co-expression with the rate-limiting enzyme in dopamine synthesis, TH, (Supplementary Figure S4Ai-ii,Bi-ii and using confocal microscopy, Supplementary Figures S3C), with > 85% of cells co-expressing TH+ and GFP+, as previously described for this reporter cell line ^5^. GFP+ neurons commonly clustered at the periphery of the graft with significant GFP-areas within the graft core (Figure 2A-B), as previously observed for both fetal and PSC-derived VM grafts. Volumetric analysis confirmed that GDNF significantly increased graft volume at 6 months (Graft: 7.04 ± 0.96 mm^3^ and Graft + GDNF: 13.33 ± 1.72 mm^3^) (Figure 2C). Both GFP+ (Graft: 5268 ± 654; Graft + GDNF: 9476 ± 1462) and TH+ (Graft: 4092 ± 602; Graft + GDNF: 7986 ± 1375) neuron numbers commensurately increased with graft volume in the presence of GDNF, such that DA cell density was unchanged (Figures 2D and 2E), indicating that GDNF promoted survival but not differentiation of DA neurons. Consistent with previous reports, TH+ DA neurons accounted for ~1% of the total graft (or ~5% of the 100,000 implanted cells) at 6 months ^3–5, 26–28^, and increased 2-fold (to ~9.5% of implanted cells) in the presence of GDNF. As an index of GDNF-induced maturation of the graft, assessment of the proportion of GFP+ (PITX3) cells that co-expressed TH revealed a subtle but not significant increase (Cells 79.29% + 9.0, Cells + GDNF 91.5% + 7.3, p=0.3, data not shown). Interestingly, the total GFP-population also increased concomitantly, suggesting that GDNF also influenced non-DA cells within the grafts (Figure 2F). Note, AAV-mCherry had no impact on motor function (data not shown) or the neural composition of the graft (inclusive of TH+HNA+ neurons, NeuN+HNA+ neurons, APC+HNA+ mature oligodendrocyte and GFAP+ astrocyte density, Supplementary Figure 3D-G), compared to grafts of cells alone and were not further analysed.

**Figure 3:**
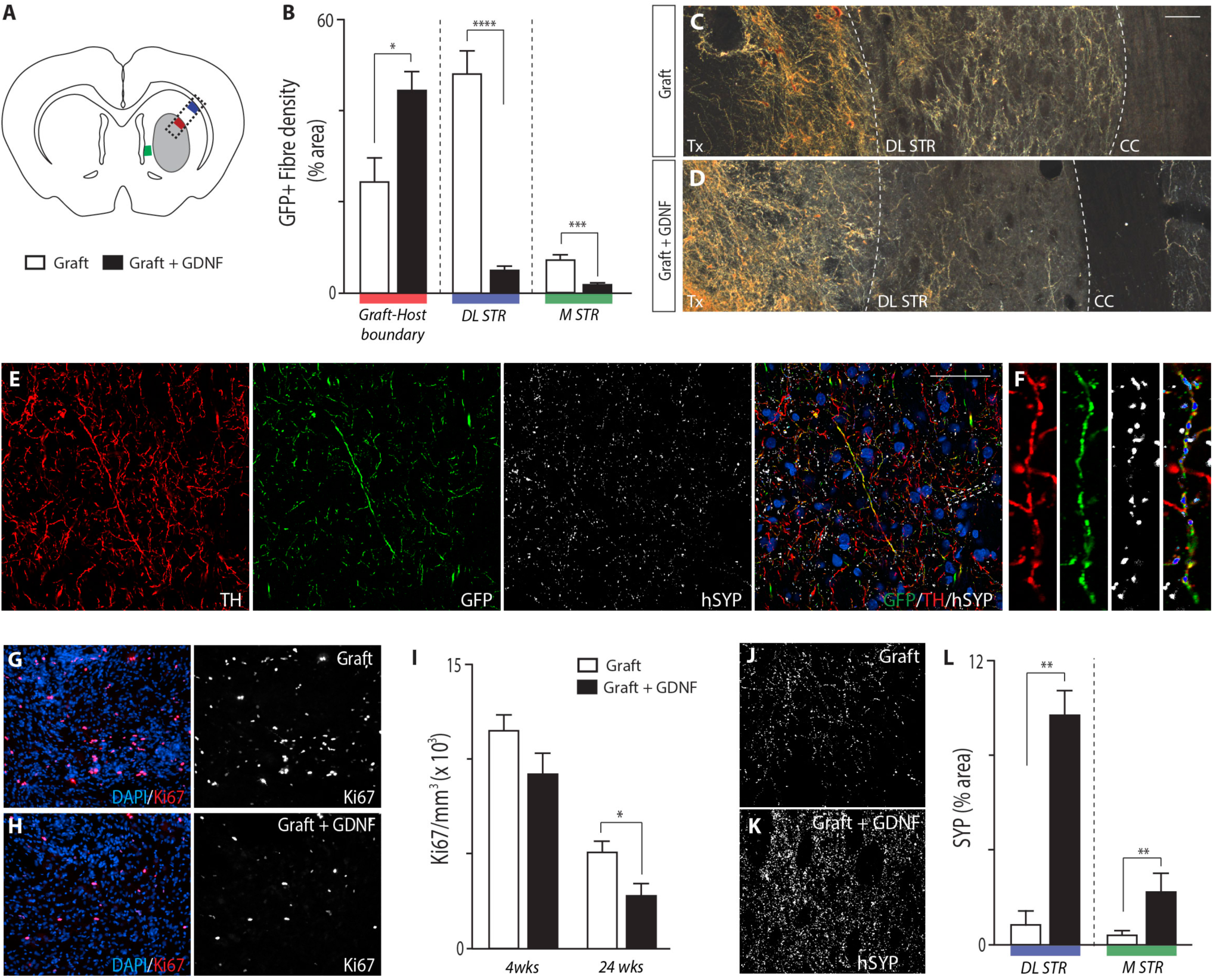
Early exposure of human PSC-derived DA transplants to GDNF impedes axonal sprouting and integration into the host brain. (A) Schematic showing sampling sites for estimates of graft-derived fiber density, at the graft-host border (red) and within the target dorsolateral striatum (blue), and region of interest depicted in C-D (dotted line). (B) Transplants in the presence of GDNF showed increased GFP+ DA innervation density at the border of the graft, but significant reduction in their capacity to innervate the host dorsolateral striatum. Data represents mean ± SEM, Students t-test (n = 5 Graft alone; n = 6 Graft+GDNF). (C-D) Representative photomicrographs illustrating the increase in graft-derived GFP+ innervation within the transplant, and decreased innervation in the striatum in animals receiving grafts in the presence (D), compared to absence (C), of GDNF. Tx, transplant; DL STR, dorsolateral striatum; CC, corpus callosum. Scale bar, 100µm. (E) Transplants were capable of forming DA synaptic connections within the host striatum, as depicted by the presence of TH+, GFP+ and hSYP co-labelling within the representative Graft+GDNF brain. Scale bar, 50 µm. (F) Example of hSYP-immunoreactive puncta along a graft derived, DA (GFP+/TH+) neurite within the dorsolateral striatum (magnification of boxed area in E). (G-H) Photomicrographs of Ki67+ proliferative cells within a graft, and in the presence of GDNF, at 24 weeks after transplantation. (I) The increase in graft size, observed in the presence of GDNF, was not due to an increase in proliferation as seen by Ki67+ analysis within the grafts at either 4 or 24 weeks post-transplantation. (J-K) Representative images, and quantitative assessment of hSYP puncta within the host striatum animals receiving a VM graft in the absence (J) and presence (K) of GDNF. (L) GDNF increased hSYP puncta density within the dorsolateral and medial striatum (DL STR and M STR, respectively), indicative of synaptic maturation and integration of the graft.

To assess the competence of transplanted DA progenitors to mature into correctly specified A9 or A10 DA subtypes we assessed Calbindin (CALB) or GIRK2 immunoreactivity within GFP+ DA neurons (Figures 2H-I). Specification of GFP+/GIRK2+ or GFP+/CALB+ was constant regardless of GDNF expression. In contrast the percentage of GFP+/GIRK2+/CALB+ co-expressing neurons significantly increased in Graft + GDNF compared to Graft animals (21.17 ± 0.97% and 41.23 ± 6.24%, respectively), such that only 16% of grafted GFP+ cells failed to adopt the A9/A10 fate in the presence of GDNF, compared to 31% of GFP+ cells in the absence of GDNF (Figure 2H, green bar). In addition, GIRK2 expressing cells within the grafts (GFP+/GIRK2+ and GFP+/GIRK+/CALB+) were significantly hypertrophied in the presence of GDNF (Figures 2J-2K), findings that collectively suggest GDNF promoted the maturation of DA neurons.

The integration of grafted DA neurons into the host tissue was assessed using GFP innervation density at the graft-host boundary, as well as within the dorsolateral striatum (DL STR), the canonical target region for the A9 DA neurons responsible for motor function and the medial striatum (Figure 3A). Reflective of the increase in GFP+ cells within the grafts, GFP+ fiber density at the graft-host boundary was significantly increased in the presence of GDNF (Graft: 24.34 ± 5.17%; Graft + GDNF: 44.05 ± 10.65% area covered by GFP+ immunoreactive pixels) (Figures 3B-D). Surprisingly, GFP+ innervation density within the dorsolateral striatum was significantly diminished in the presence of GDNF (Graft: 47.25 ± 5.39%; Graft + GDNF: 4.8 ± 0.93%). To determine whether the reduced innervation was a consequence of excessively high GDNF protein levels, we assessed GFP+ fiber density within the medial striatum, where protein levels were notably lower (Supplementary Figure 2A). Similar to the dorsolateral striatum, GFP+ fibre density was significantly reduced (Figure 3B). Despite poor host innervation from grafts in the presence of GDNF, those GFP+ fibers within the dorsolateral striatum remained capable of forming synapses, as revealed by the presence of human-specific synaptophysin (hSYP) immunoreactivity along GFP+ fibers (Figure 3E-F).

The surprising reduction in DA innervation of the host striatum by grafted cells exposed to GDNF led us to delve further into understanding the maturation, differentiation and plasticity of these transplants that were capable of improving motor function. Quantitative assessment of KI67+ proliferative progenitors within the grafts at 4 and 24 weeks after transplantation revealed a progressive and subsequently significant decrease (Figure 3G-I), indicating the pro-maturation (and not proliferative) effect of GDNF on the graft. Confirming these maturation findings, the density of NeuN+ cells within GDNF exposed grafts was significantly increased (Cells: 41.6% + 1.9 NeuN+HNA+ immunoreactive cells, and Cells+GDNF: 58.0% ± 3.0, p=0.0009, Figure 2G). Total 5HT+ serotonergic neurons, a neuronal population capable of dopamine uptake and release, influencing motor function and contributing to graft-induced dyskinesias ^29, 30^, were also significantly elevated 3-fold in GDNF exposed grafts (5HT+HNA+Cells: 502 ± 11 and Cells+GDNF: 1420 ± 206). Other neuronal populations (inclusive of GAD67+ GABAergic, DBH+ noradrenergic, ChAT+ cholinergic) were notably sparse in the grafts and not different between Graft and Graft+GDNF. Reflective of the increased neuronal maturation, the density of graft-derived human synaptophysin (hSYP+) synapses were also more abundant in the presence of GDNF, within both the dorsolateral (Graft: 1.24 ± 0.52%; Graft + GDNF: 8.52 ± 1.69% hSYP+ area) and medial striatum, (Figure 3J-L). Together, these data suggest that whilst GDNF induced graft maturation and synaptogenesis, effects were also observed in non-DA neurons, inclusive of undesirable serotonergic cells that have the capacity to influence motor functions.

### Delayed GDNF delivery promotes recovery of motor function without increasing cell survival

As the DA neurons only accounted for a fraction of the graft, a common observation for human PSC-derived VM progenitor grafts ^3-5, 27, 28^, and that the delivery of GDNF prior to cell transplantation negatively impacted on the ability of the grafted DA neurons to extend axonal projections outside of the graft core to innervate the host striatum, we looked to employ two major refinements to a subsequent study design. First, we employed an LMX1A-eGFP reporter ESC line to enable selective isolation of VM progenitors prior to transplantation, based upon recent findings that grafts of LMX1A-GFP+ progenitors enriched for DA neurons 3-fold ^31^. Second, we altered the timing of GDNF delivery to 3 weeks after transplantation, such that the DA-enriched grafts were exposed to GDNF after a period that would influence survival, addressing instead the capacity for the neurotrophic factor to modulate axonal plasticity. These modifications were deemed an advancement of the study and not intended for comparative assessment of grafts exposed to GDNF from the outset (assessed in figure 1-3).

As previously demonstrated ^2^, we confirmed the ability of the LMX1A-eGFP H9 hESC line to differentiate into VM progenitors, with the majority of cells (>85%) co-expressing GFP, OTX2 and FOXA2+ at 15DIV (Figure 4A). On the day of transplantation (21DIV) the cultures yielded a high proportion of immature GFP+ (LMX1A)/NURR1+/TH+ progenitors (Figure 4B) that could be FACS-purified (>70% eGFP+, Figure 4C) and transplanted directly into the denervated striatum.

**Figure 4:**
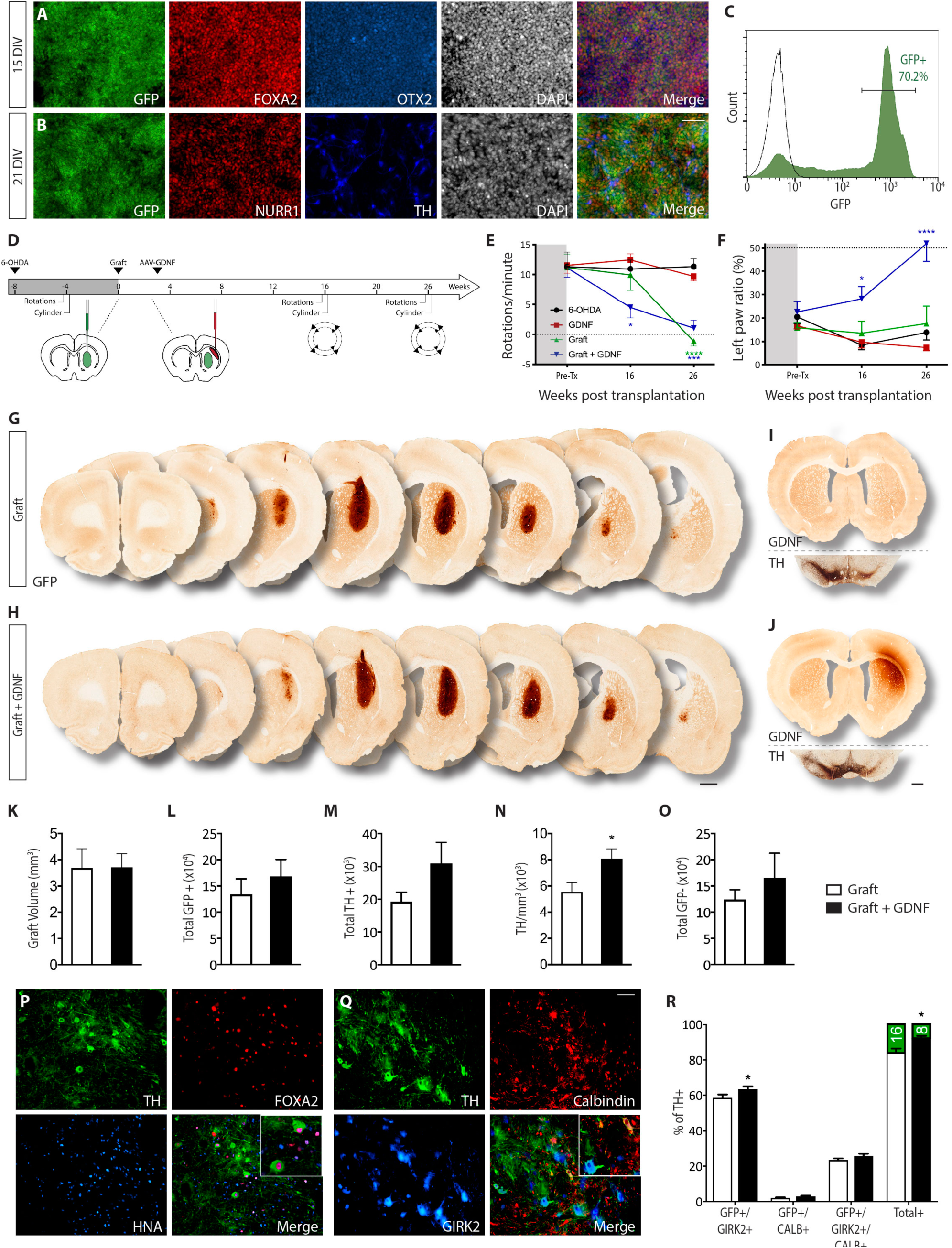
Delayed exposure of hPSC-derived DA grafts to GDNF enhances functional recovery in Parkinsonian rats, independent of cell survival. (A-B) Differentiation of LMX1A-eGFP hESC into VM progenitors was confirmed by the high co-expression of GFP, FOXA2 and OTX2 by 15DIV (A) as well as GFP, NURR1 and TH at the time of transplantation (21DIV) (B). Scale bar, 50µm. (C) Fluorescent activated cell sorting plot illustrating the isolation of LMX1A-eGFP expressing VM progenitors from differentiating cultures at 20DIV for transplantation. (D) Schematic overview of the study design inclusive of behavioural testing, cell transplantation and AAV injections. (E-F) All grafted animals showed restoration of amphetamine-induced rotational asymmetry at 26 weeks after grafting (E), yet only rats receiving transplants in the presence of GDNF showed significant improvements in the use of the impaired (left) paw in the cylinder test (F). Lesion, n = 9; GDNF, n = 8; Graft, n = 7 and Graft + GDNF, n = 7. Two-way ANOVA against 6-OHDA with Dunnett correction for multiple comparisons. (G-H) Representative overview of a hESC-derived VM progenitor transplant in the absence (G) and presence (H) of AAV-GDNF. Grafts were immunostained for GFP to specifically identify LMX1A-eGFP expressing graft deposits. Scale bar, 1mm. (I) Coronal section illustrating the absence of GDNF expression in the adult striatum, and reduction in TH-ir in the lesioned midbrain of the rat brain depicted in (G). (J) GDNF immunohistochemistry confirmed targeted, long-term delivery of the AAV-GDNF into the dorsolateral striatum, and overlying cortex. TH immunohistochemistry confirmed MFB DA lesioning. Images are from the same brain depicted in (H). (K-N) Graphs highlighting that delayed GDNF delivery had no significant effect on graft volume (K), the survival of GFP+ DA progenitors (L) or TH+ DA neurons (M) but has a modest effect on TH+ cell density (O) GDNF also had no effect on the total number of non-DA (HNA+/TH-) cells within the graft. (P) The ability of LMX1A+ grafted progenitors within the graft to mature into DA neurons was confirmed by the co-expression of TH, FOXA2 and HNA. (Q) High proportions of transplanted DA neurons (confirmed by TH-immunoreactivity) co-expressed GIRK2 and/or Calbindin (CALB), indicative of A9 and A10-like specification. Scale bar (P,Q): 50µm. (R) Quantification of TH+ neurons that co-express GIRK2 and/or CALB. For panels K-O,R, data presented as mean ± SEM, Students t-test (n = 7 rats/group).

Amphetamine induced rotation and cylinder tests were conducted periodically before and after grafting to validate the functionality of LMX1A-GFP derived DA transplants ± GDNF (Figure 4D). 6-OHDA-induced motor deficits were again stable for 26 weeks in 6-OHDA lesion control animals in the absence and presence of GDNF (Figure 4E, black and red lines, respectively). Complete abolishment of amphetamine-induced rotational asymmetry was observed 26 weeks after grafting in animals receiving transplants (with or without GDNF, blue and green lines, respectively). However, recovery was notably accelerated in animals with combined DA grafts and GDNF (Figure 4E, blue line). In the cylinder test, only animals receiving combined graft and GDNF treatment showed significant improvement in contralateral forepaw use (Graft + GDNF: 51.94 ± 7.66%, blue line; Graft: 17.72 ± 7.40%, green line; GDNF alone: 7.49 ± 1.51%, red line) (Figure 4F).

At 26 weeks post-transplantation, surviving grafts, identified as dense GFP immunoreactive deposits, were observed in all animals (Figures 4G and 4H). Sustained GDNF expression, restricted to the dorsolateral striatum and overlying cortex, was confirmed only in animals that received AAV-GDNF (Figures 4I and 4J). 6-OHDA lesions consistently ablated TH+ neurons within the substantia nigra, while leaving ventral tegmental area DA neurons relatively spared (Figures 4I and 4J). LMX1A+ grafts were confirmed to be rich in midbrain dopaminergic neurons by 6 months, as indicated by the co-expression of TH, FOXA2 and human nuclear antigen (HNA), Figure 4P. As anticipated, quantitative assessment revealed no difference in graft volume (Figure 4K), GFP+ cells (Figure 4L) or TH+ DA neuron number between the two groups (Graft: 19,810 ± 2,828; Graft + GDNF: 30,381 ± 6,513, P=0.153, Figure 4M), highlighting the absence of a cell survival effect. However, total yields of TH+ DA neurons within both Graft and Graft + GDNF were substantially elevated following sorting, accounting for 20-30% of the 100,000 implanted cells, and an estimated 10% of total cells within the transplant (TH+/HNA+). The high proportion of DA neurons within the graft could be most evidently seen by comparative assessment of PSA-NCAM (total graft), GFP (VM progenitors and DA neurons) and TH+ (DA neurons) (Figure S3), reflecting the efficacy of the selective isolation and transplantation of appropriately patterned VM-specific LMX1A-GFP+ progenitors. Graft exposure to GDNF had a modest but significant impact on TH+ density (8095 ± 730/mm^3^) compared to grafts of DA progenitors alone (5,555 ± 699/mm^3^) (Figure 4N). Assessment of the non-dopaminergic fraction of the graft (TH-/HNA+ cells) revealed no change in cell number, indicative of the absence of a GDNF effect on survival and/or proliferation (Figure 4O). No difference was observed in the proportion of NeuN+/HNA+ neurons, APC+/HNA+ mature oligodendrocytes or the density of GFAP+ astrocytes within Cell or Cell+GDNF grafts, data not shown. Furthermore, 5HT+ neurons were notably sparse within the grafts, in the presence or absence of GDNF – a positive consequence of LMX1A+ VM progenitor selection prior to grafting (de Luzy et al., 2019).

Co-expression of TH, together with GIRK2 and/or Calbindin again confirmed the ability of DA neurons within the grafts to mature into subtype specific populations with similar proportions in the presence or absence of GDNF (Figures 4Q and 4R). Small but significant increases in the overall proportion of Total GIRK2 and/or CALB specified DA neurons were seen in animals receiving GDNF (Graft: 84.44 ± 1.96%; Graft + GDNF: 92.24 ± 1.41%), indicative of improved and/or accelerated maturation of the grafts at 26 weeks (Figure 4R).

### Delayed delivery of GDNF modulates appropriate innervation of the host targets from human stem cell derived DA neurons

Given the robust number of TH+ neurons present within the grafts, we next assessed the modulatory properties of GDNF on graft-derived DA innervation within key target nuclei of the midbrain dopamine system. As GFP (LMX1A) was not selectively indicative of DA identity, TH-immunoreactivity was used to quantify the striatal innervation density of DAergic fiber terminals. Important for the quantitative assessment of TH+ DA fibers emanating from the graft was the need to confirm robust ablation of the host DA system within the ipsilateral striatum and overlying cortex, as observed in 6-OHDA lesioned animals (Figure 5A). In contrast, notable TH immunoreactivity was observed along the rostrocaudal axis ipsilateral to the graft in animals that received transplants (±GDNF) (Figures 5B and 5C). Delayed exposure of the graft to GDNF (three weeks post-transplantation) resulted in significant increases in TH+ fiber density within multiple intra-and extra-striatal nuclei, including the overlying cingulate cortex (Cing. CTX), perirhinal cortex (Peri. CTX), dorsolateral striatum (DL STR) and ventrolateral striatum (VL STR) (Figures 5B-D).

**Figure 5:**
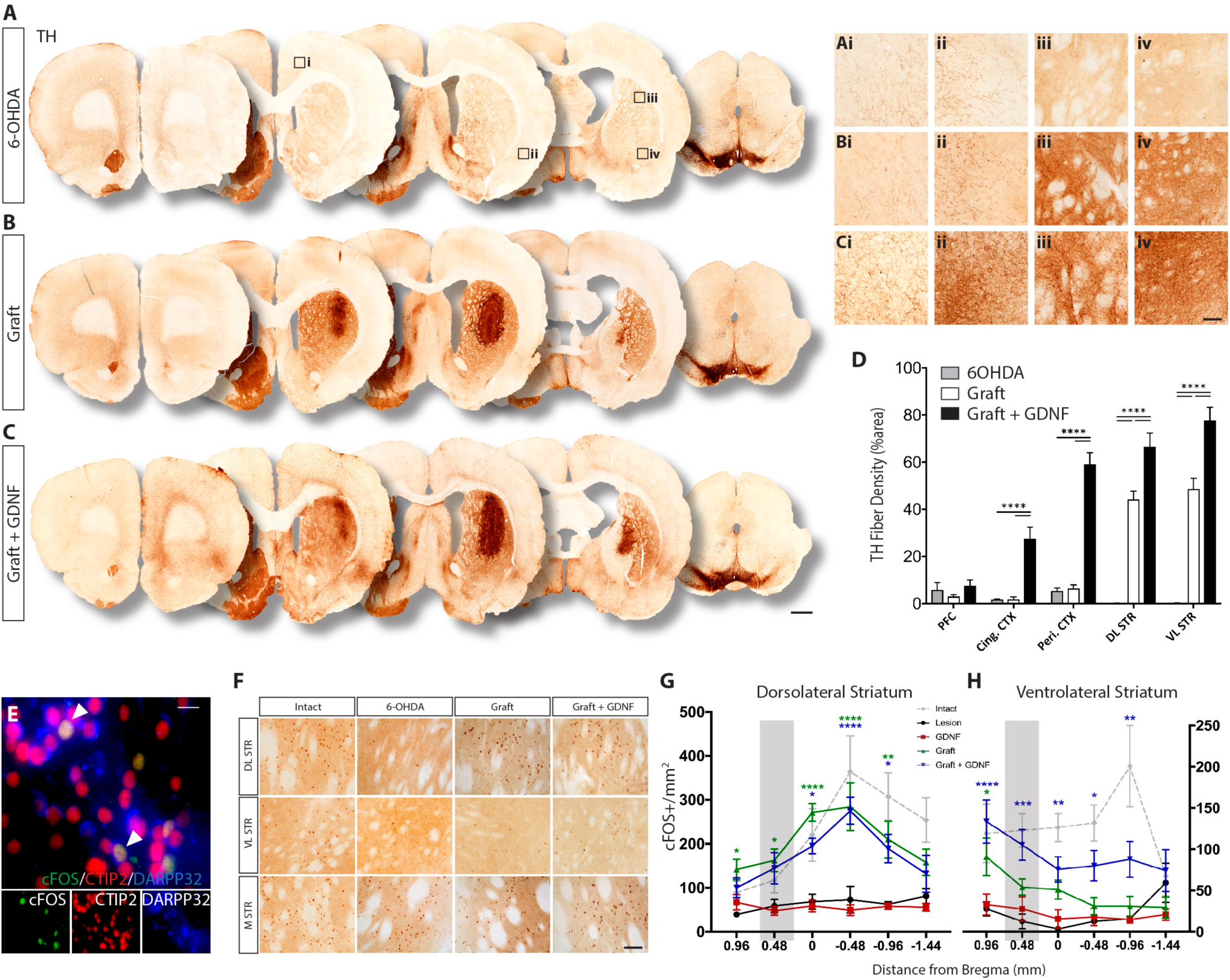
Delayed exposure of the VM graft to GDNF promoted dopaminergic innervation of the host tissue, activation of striatal neurons and upregulated dopamine biosynthesis. (A-C) Photomontages illustrating the density of TH+ fibers within the rat brain following 6-OHDA lesioning (A) and after cell transplantation in the absence (B) and presence of AAV-GDNF (C). (Ai-Ci) High power images from (A-C) illustrating the density of TH+ fibers in the motor cortex, (Aii-Cii) perirhinal cortex, (Aiii-Ciii) dorsolateral striatum and (Aiv-Civ) ventrolateral striatum. Note the increase in TH+ fibers in striatal and extrastriatal regions of animals receiving grafts in the presence of GDNF. Scale bar, 1mm (A-C) and 100μm (insets). (D) Quantification of TH fiber density within the host tissue of 6-OHDA lesioned rats with and without cell grafts and AAV-GDNF. Two-way ANOVA, Tukey correction for multiple comparisons. 6-OHDA Lesion, n= 9; Graft, n= 7; Graft + GDNF, n=7 rats. (E) Activation of host striatal neurons was confirmed by co-expression of c-FOS, CTIP and DARPP-32. Scale bar, 40μm. (F) Representative photomicrographs of c-FOS+ cells within the striatum of Intact and 6-OHDA lesioned rats ± Grafts ± AAV-GDNF. (G) Quantification of c-FOS+ cells within the host brain revealed significant increases in activated cells in the dorsolateral striatum of grafted animals, irrespective of GDNF expression, compared to ungrafted rats (Lesion ± GDNF). (H) Rats receiving grafts in the presence of GDNF showed significant increases in cFOS activation in the ventrolateral striatum. For panels G-H: Panels G-H, data represents Mean ± SEM. Two-way ANOVA against 6-OHDA, Dunnett correction for multiple comparisons. Intact, n = 6; Lesion, n = 9; GDNF, n = 8; Graft, n= 7; Graft + GDNF, n=7. Grey bar represents the site of transplantation (0.5mm rostral of bregma). Abbreviations: PFC, prefrontal cortex; cing. CTX, cingulate cortex; peri. CTX, perirhinal cortex; DL STR, dorsolateral striatum; VL STR, ventrolateral striatum.

Acute upregulation of the intermediate early response gene, c-FOS, in medium spiny neurons (MSN) of the striatum following amphetamine administration has been shown to be modulated by DA release and subsequent receptor activity, thereby providing an indirect measure of DA signalling ^32^. To facilitate c-FOS analysis, all animals were injected with d-amphetamine one hour prior to perfusion. c-FOS+ activation, was validated in striatal MSNs by CTIP2+/DARPP-32+ co-expression (Figure 5E), with quantitative assessments performed within the medial, dorsolateral and ventrolateral striatum (Figured 5F-H and Supplementary Figure S5). As anticipated, 6-OHDA lesioning (in the absence of a graft) reduced c-FOS expression within both the dorsolateral and ventrolateral striatum, in comparison to the contralateral hemisphere and intact control animals (Figure 5F-H and Supplementary Figure 4C-D). Lesioning had no effect on c-FOS+ cells in the medial striatum (Figure 5F). Grafts (± GDNF) increased c-FOS+ density along the rostrocaudal axis of the dorsolateral striatum, an effect particularly pronounced immediately caudal to the site of graft implantation, and not significantly different to the intact control animals (Figure 5G). Within the ventrolateral striatum, only grafts in the presence of GDNF showed a significant increase in c-FOS expression relative to the lesion controls (Figure 5H), that was not significantly different to the intact brain. Within the caudal regions of the ventrolateral striatum (0.5 to 1.5mm caudal to bregma), density of cFOS+ was significantly elevated in animals receiving grafts in the presence, compared to absence of GDNF (Supplementary Figure S5D).

### GDNF restores dopamine levels and modulates graft-specific gene expression

Following the establishment of GDNF’s modulatory effect on DA neuronal survival and innervation patterns, we next investigated the effect of GDNF on gene expression and DA metabolism within the graft. A cohort of mice were transplanted with FACS-purified LMX1A+ DA progenitors (± AAV-GDNF 3 weeks after transplantation), with tissue isolated at 6 months for either transcriptional profiling or high-performance liquid chromatography (HPLC), (Figure 6A-B). Mice were confirmed to have comparable, viable grafts at 6 months to those observed in rats (Supplementary Figure 6A). To identify gene changes within the graft, we performed species-specific RNA-seq analysis on host striatal tissue, containing the VM grafts ± GDNF, using previously described methods^33^. Whole genome hierarchical analysis revealed distinct differences in gene expression between the two graft groups, with 240 genes significantly upregulated and 40 genes downregulated in the Graft + GDNF compared to Graft alone (wFDR<0.05, Figure 6C-D). GDNF-upregulated genes were associated with gene ontology biological processes including cell-cell signalling, synaptic signalling, cell communication and secretion (Figure 6E).

**Figure 6:**
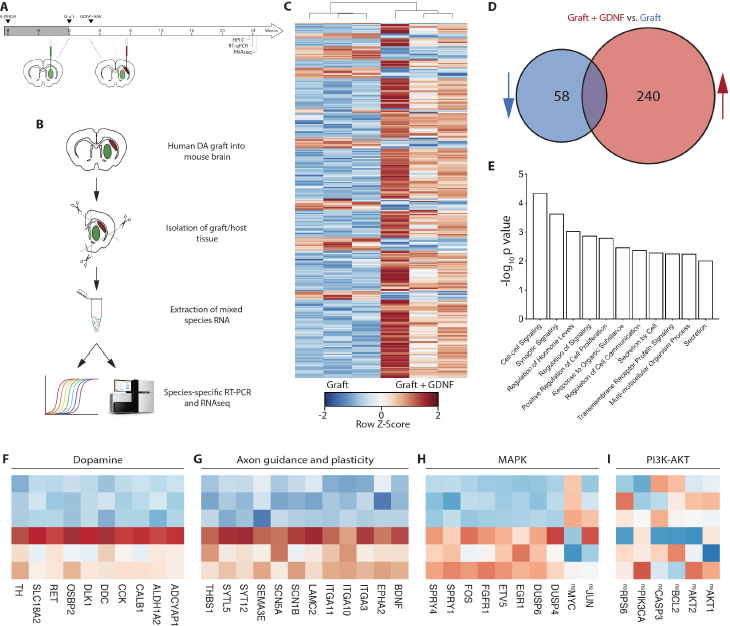
Transcriptional profiling of VM grafts in the presence and absence of GDNF identified known and novel DA homeostatic, synaptogenesis and axonal plasticity-related genes. (A) Schematic overview of experiments designed to probe transcriptomic and DA metabolic modulation (± GDNF) (B) RNA was isolated from grafted mice (± GDNF) and analysed by both RT-qPCR and RNAseq. (C) Unbiased hierarchical clustering analysis of gene expression changes in Graft and Graft + GDNF tissue samples, (n=3/group). (E) Gene ontology analysis clustering of upregulated genes in Grafts exposed to GDNF (<0.01 FDRl ‘Synaptic signalling’ includes related GO terms with equal ‘Anterograde trans-synaptic signalling’, ‘Trans-synaptic signalling’ and ‘Chemical synaptic transmission’). Note, ‘ns’ preceding gene names indicates those ‘not significantly’ changed. (F-I) Heat map of selected genes related to dopamine (F), axon guidance and plasticity (G), MAPK signalling (H) and PI3K-AKT signalling (I).

Selective analysis of the upregulated genes from the RNAseq revealed many with known roles in dopamine development and homeostasis such as TH, SLC18A2/VMAT, DDC/AADC and ALDH1A2 (Figure 6F). The non-canonical Notch ligand DLK1, known to influence VM DA neurogenesis and DAT expression in development ^34, 35^, be regulated by GDNF ^36^, and recently identified as a predictive marker of good human PSC-derived DA graft outcomes (TH+ cell number and innervation density) ^26, 37^ was also upregulated. Supporting the influence of GDNF on graft plasticity and motor function, upregulation of numerous synaptic (SYTL5, SYT12), axon guidance (BDNF, EPHA2, SEM3E) and adhesion (LAMC2, ITGA11/10/3) genes were also observed (Figure 6G).

A number of the transcriptionally upregulated genes identified by RNAseq were verified by qPCR, using human-specific primers to target the graft, and not host, transcript (Supplementary Figure 5C). These RNAseq/qPCR findings provide the first detailed transcriptional profiling of hPSC-derived grafts in response to extrinsic modulation, giving insight into the underlying genetic changes that drive anatomical rearrangement and physiological outcomes, to here, improve the functional integration of DA-enriched grafts. Such profiling analysis also provide new insight into graft responses. By way of example, here we show cholecystokinin (CCK), a neuropeptide known for modulating DA release within midbrain dopamine pathways, including in fetal VM transplants ^38^, was significantly upregulated in response to GDNF (Figure 6F, Supplementary Figure 5C). To confirm the observed increase at the transcriptional level, we examined the expression and localisation of CCK protein. We showed VM grafts in the presence of GDNF had 2-fold more CCK+ neurons, with 34.6% co-expressing TH, suggesting that GDNF regulates CCK expression and potentially modulates DA transmission within the grafted brain. While CCK cell counts were low, this reflects the challenges in visualising this population at a protein level, noting that the peptide is rapidly translocated to the nerve terminal^39^. With CCK expressed within most midbrain DA neurons^40^, further studies involving blocking of axonal transport, will be required to reveal the true impact of GDNF of this population. Protein-protein interaction network analysis^41^ of differentially expressed genes demonstrated the upregulation of genes associated with neural development, synaptic transmission, adhesion and GDNF activity, further confirming our observations within maturing DA grafts (Supplementary Figure S7). Further studies will be required to fully elucidate the role/s of CKK, and other novel GDNF-downstream genes on DA survival, plasticity and function.

We additionally assessed genes associated with downstream GDNF signalling. Within DA neurons, GDNF/GFRα1 binding triggers signalling via a complex with the receptor tyrosine kinase Ret, which in turn modulates activation of downstream PI3K-AKT and MAPK-ERK pathways, that activate transcription of genes involved in survival and/or plasticity ^42^. In addition to significantly elevated Ret expression (Figure 6F), we observed upregulation of a number of MAPK-associated genes with known roles in plasticity, including EGR1 and SPRY1 (two predictive genes recently linked to high dopamine neuron yields within in grafts^26^), but no change in MAPK-associated genes linked to cell migration and proliferation (JUN, MYC), Figure 6H. In contrast, genes associated with the PI3K-AKT pathway (e.g. AKT1, AKT2 and PIK3CA) were not significantly changed (Figure 6I), suggesting that GDNF mediates the plasticity of human DA neurons via the MAPK-ERK, and not PI3K-AKT, pathway.

These findings were confirmed by assessing the expression of phosphorylated ribosomal protein S6 (pS6), an indicator of the upstream activation of the neuroprotective pathway Akt/mTOR pathway, as well as phosphorylated ERK (pERK), specifically in graft-derived (PITX3-GFP+) DA neurons. Assessment of the host midbrain (ipsilateral and contralateral to the AAV-GDNF intrastriatal delivery) verified the capacity to measure changes in pERK and pS6, as revealed by increased labelling within TH+ neurons (Figure 7A-B). Confirming the transcriptional findings from grafted animals, a significant increase in the proportion of DA neurons co-expressing pERK (%pERK+GFP+/Total GFP+) was observed in grafts exposed to late/delayed GDNF (Cells: 21% ± 5; Cells+GDNF: 48% ± 6), yet no change was observed the proportion of pS6-immunoreactive DA neurons, Figure 7D,G.

**Figure 7:**
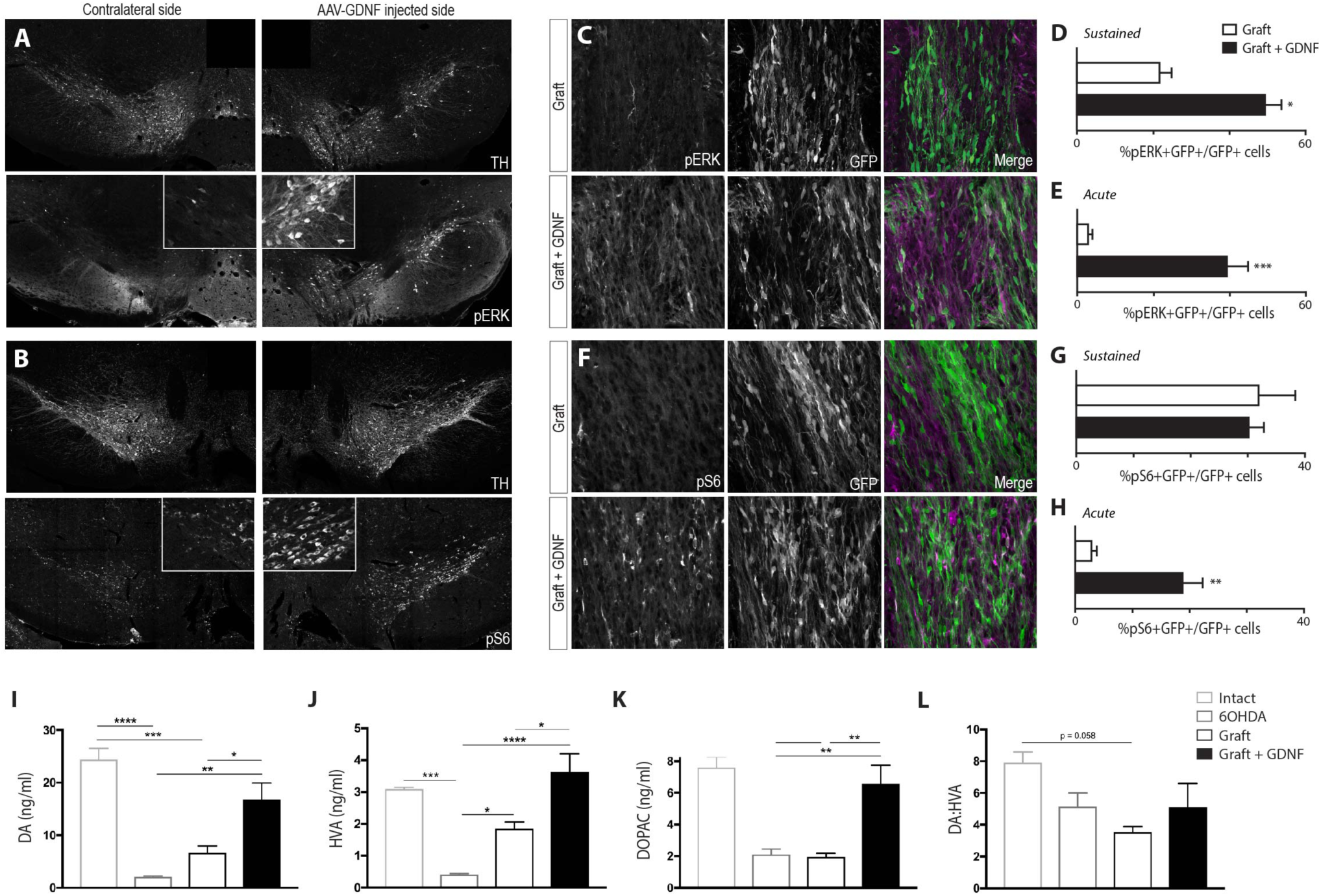
GDNF overexpression temporally regulates downstream pERK and pS6 signalling in grafted dopamine neurons. (A-B) Assessment of the host midbrain (ipsilateral and contralateral to the AAV-GDNF intrastriatal delivery) verifying the capacity to measure changes in (A) pERK and (B) pS6 within host, TH+ DA neurons. (C) Representative images showing pERK co-localised with GFP+ (PITX3) DA cells in grafts in the absence or presence of ‘Acute’ GDNF exposure. (D-E) Proportion of GFP+ DA neurons co-expressing pERK after ‘Sustained’ (D) and ‘Acute’ (E) exposure to AAV-delivered GDNF. (F) Representative images showing pS6 co-localised in GFP+ DA cells within grafts in the absence or presence of ‘Acute’ GDNF exposure. (G-H) Proportion of GFP+ DA neurons co-expressing pS6 after ‘Sustained’ (G) and ‘Acute’ (H) exposure to AAV-delivered GDNF. (I-K) HPLC confirmed a significant reduction in dopamine (I) and DA metabolites, HVA (J) and DOPAC (K), in the striatum of lesioned (compared to intact) mice, an effect that could be partially restored by hESC-derived DA grafts, and fully restored when transplants were exposed to GDNF. (L) Reflective of commensurate changes in DA and it’s metabolites, dopamine turnover (DA:HVA) remained unchanged across lesion and grafted animals. For panels D,E,G,H,I-L, Data represents mean ± SEM, n=4-6 mice/group. (D,E,G,H) Students t-test; (I-L) One-way ANOVA with Tukey correction for multiple comparisons.

Recognising that assessment of PI3K-AKT and MAPK-ERK was performed on grafts that had been exposed for many months to GDNF, and the potential for modulation of downstream target genes over time, we performed an additional cohort of transplantation (± ‘late GDNF’) to assess acute responsiveness of the grafts, within days of exposure, to GDNF protein. Consistent with transcriptional and protein changes in response to sustained GDNF exposure, acute exposure to GDNF induced a 10-fold increase in pERK-expressing DA neurons, compared to within grafts not exposed to GDNF (Figure 7C,E). In contrast to sustained GDNF, acute exposure significantly increased the proportion of DA neurons co-expressing pS6 (Figure 7F,H), suggestive of early dual GDNF roles in survival and plasticity, with ongoing contributions to graft plasticity at protracted periods after transplantation.

Finally, we utilised HPLC to demonstrate that striatal DA metabolism was elevated in animals receiving grafts in the presence of GDNF. Whilst not a direct assessment of physiological DA release (due to analysis of brain homogenates), comparative assessments of DA levels, and metabolites, could be made between groups. Importantly, a >90% reduction in striatal dopamine levels in 6-OHDA lesioned animals confirmed ablation of the host midbrain dopaminergic system, (6-OHDA: 2.0 ± 0.2pmol/mg of wet tissue; Intact: 24.4 ± 2.2pmol/mg) (Figure 6J). Reflective of the improvement in rotational asymmetry at 24 weeks, grafts in the absence or presence of GDNF showed significant elevations in DA levels (6.7 ± 1.3pmol/mg and 16.8 ± 3.1pmol/mg, respectively) (Figure 6J), as well as the DA metabolites HVA (1.9 ± 0.2pmol/mg and 3.6 ± 0.6pmol/mg) and DOPAC (1.9 ± 0.3pmol/mg and 6.6 ± 1.1pmol/mg), compared to 6-OHDA lesion alone (HVA: 0.4 ± 0.03pmol/mg; DOPAC: 2.1 ± 0.3pmol/mg) (Figures 6K and 6L). However, only in the presence of GDNF were DA, DA metabolites and overall DA metabolism (ratio of DA to HVA) restored to levels not significantly different from the intact brain, reflective of restoration of dopamine biosynthesis and function in these animals (Figures 6J-M). These increases in DA levels seen in graft tissue exposed to GDNF was supported by elevated expression of genes associated with DA synthesis gene, including TH and AADC (Figure 6F, Supplementary 5C).

## Discussion

This study provides the first evidence of modulating the survival and plasticity of human pluripotent stem cell-derived dopaminergic transplants in the Parkinsonian brain. Employment of unique GFP-expressing stem cell reporter lines enabled precise tracking of the DA contribution to the grafts (PITX3-eGFP), as well as a 10-fold enrichment of DA neurons (by FACS isolation LMX1A-GFP progenitors) within the transplants. Utilizing these tools, we were able to demonstrate that the timing of onset for sustained GDNF delivery directly impacted on the mechanisms by which improvements in functional recovery occurred. While all grafted animals showed restoration of amphetamine-induced motor asymmetry by 6 months, reflective of the relatively low number of DA neurons required for correction of this gross motor deficit (estimated to be as few as 1000 human DA neurons ^6^), examination of non-pharmacologically-induced sensorimotor tasks revealed distinct benefits of sustained GDNF expression.

We demonstrate that implantation of human PSC-derived VM progenitors into an already GDNF-enriched environment promoted graft survival, yet without selectivity for the DA population. More noteworthy however was the negative impact of early GDNF delivery on the capacity of the graft to innervate the host tissue, such that the majority of graft-derived DA fiber growth was restricted to the graft core. Within these animals we demonstrate increased neuronal differentiation and striatal innervation from non-DA neurons, inclusive of 5HT neurons, that likely contribute to the functional recovery observed, together with the increased maturation (GIRK/CALB specification) of TH+ DA neurons within these grafts. Further studies, such as selective silencing of DA cells within grafts^43^, will be required to elucidate the relative contribution of TH+ DA neuron verses other neuronal populations to behavioural responses. In contrast, delayed exposure of the graft to GDNF had no impact on cell survival, yet significantly increased graft-derived DA reinnervation of the host tissue, elevated striatal DA levels (to levels not different from the intact brain), and consequently, enhanced activation of postsynaptic striatal medium spiny neurons.

We demonstrate that these effects observed in the present study are mediated through acute PI3K-AKT and sustained MAPK signalling, downstream of GDNF-receptor activation. Interestingly, while increased activation of striatal neurons (the primary output of the transplanted DA neurons) was shown in the dorsolateral striatum of animals receiving grafts in the absence and presence of GDNF, only in the presence of GDNF was cFOS activation observed in the ventrolateral striatum, a region of the basal ganglia previously shown to underpin changes in more complex sensorimotor tasks following lesioning and DA transplantation ^44, 45^, and supported by findings in patient fetal grafting studies ^46^. The importance of these findings are significant in light of former clinical trials where an estimated 15% of patients developed graft-induced dyskinesias, a phenomenon speculated to be, in part, the consequence of incomplete and uneven striatal DA reinnervation ^29, 47^.

A major contributor to the functional capacity of VM DA grafts is the relative contribution of A9 neurons^48^. Previous studies have described differential effects of GDNF on DA subpopulations, such that A10, but not A9 neurons, are protected into adulthood against apoptosis, resulting in increased cortical, but not striatal innervation ^49, 50^. By contrast, others have linked GDNF levels with the survival of A9, but not A10, DA neurons ^51^. In accordance with our previous work ^5^, here we report ~70% of TH+ cells within the graft adopted an A9-or A10 like identity, with the majority of these fate-specified cells expressing GIRK2 (alone or in combination with Calbindin), while relatively few TH+ cells expressing Calbindin alone. Such proportions are reflective of the A9 and A10 contributions within the human ventral midbrain ^52^. Surprisingly, in response to GDNF, a significant increase in TH+GIRK2+ or TH+GIRK2+CALB+ cells was observed, such that fewer than 16% of the TH+ cells remained unspecified in response to early GDNF, (and just 8% after delayed GDNF delivery). This increase in GIRK2-expressing cells in the presence of GDNF was also accompanied by increased soma size, suggestive of a role for GDNF in the maturation of human DA neurons, an effect that may contribute to improvements in complex motor tasks.

In addition to the effects on survival and plasticity of DA neurons, GDNF has been shown to influence dopamine biosynthesis through the regulation of *TH* gene expression, a response dependent on *Ret* ^53^. Unlike rodent studies where sustained GDNF induces the long term downregulation of *TH* mRNA, a compensatory response to increased DA levels from the graft resulting in impaired graft function ^23, 24^, non-human primate studies have reported maintained increases in TH+ cells and fiber density ^21, 54, 55^. For the first time, here we report the responsiveness of human dopamine neurons to chronic GDNF exposure. In alignment with non-human primates we observed increases in TH+ cells and TH+ fibers (dependent on the time of GDNF delivery) that was accompanied by significant upregulation of *TH* mRNA. We also detected elevated levels of aromatic L-amino acid decarboxylase (*AADC)*, the enzyme required for the conversion of L-DOPA into dopamine and CCK, a neuropeptide previously reported to modulate striatal dopamine levels, suggestive that GDNF is capable of regulating DA generation and release at multiple levels. Such observations were also supported by increased DA synthesis from grafts in the presence of GDNF, such that levels were not significantly different to the intact midbrain system. Noting previous pre-clinical and clinical studies observations, that developing rather than mature DA neurons are most responsive to GDNF ^21^, the maintained elevated enzyme (*TH* and *AADC*) transcript levels observed here likely reflect ongoing graft maturation, an unsurprising observation in light of the duration of human embryonic development as well as the protracted time for integration of human stem cell and fetal DA grafts in pre-clinical and clinical trials to date ^5, 25^. In this regard, extended studies, involving graft assessment at time points when human DA neurons have reached functional maturity and no longer require GDNF, will be needed to determine the effects of sustained GDNF on *TH* and *AADC* transcript levels, resultant DA biosynthesis and motor function.

GDNF expression is recognised to be tightly regulated during embryonic development, with expression levels downregulated in the adult brain, beyond periods of critical cell survival and target acquisition ^10^. Hence, it is probable and likely that closer attention to the duration of expression may be required when employing gene therapy approaches targeted at promoting neuroprotection and/or plasticity. While the present findings highlight the impact of exposing the grafted cells from either the outset of implantation or after a delayed period, the impact of sustained expression remains unknown for human stem cell-derived DA neurons. In addition to potential changes in regulatory genes of DA synthesis, further studies will be required to, for example, determine whether sustained expression may result in increased aberrant innervation of target tissue that results in unregulated synaptic neurotransmission and/or increased innervation of off-targets, as a consequence of suboptimal gene therapy targeting. The dual effects of early and late GDNF on grafted DA neurons suggest that the desired therapeutic approach may involve phasic exposure to GDNF – such that acute GDNF delivery at the time of implantation can promote survival, with a subsequent exposure within the target tissue at a defined period (potentially weeks to months) after cell transplantation. With the recent demonstration of inducible expression vectors, for the temporal control of GDNF in the intact brain^56^, this presents a feasible consideration for the future.

Added to requirements for temporal regulation is the need for greater control over gene therapy-associated protein dose. Here we report targeted injection of 1.8×10^9^ genomic copies of the AAV-GDNF into the striatum of rats. While histochemical analysis confirmed protein expression, resulting in anticipated and positive impacts on graft survival and plasticity, knowledge of absolute protein levels here remained unknown. Previous reports have demonstrated that it is in fact possible to overdose the system and impart effects on other neuronal populations ^21^ as well as negatively regulate DA levels ^23^, such that lower chronic doses may be more beneficial. Graft-derived DA innervation within the dorsolateral striatum of GDNF treated animals was reduced compared to non-GDNF animals, and similarly observed within the medial striatum, where GDNF proteins levels were notably lower, suggesting that within the current context, the presence or absence of the protein, rather than concentration, had the greatest impact on plasticity.

Added to this, dependent on the desired requirement, culture studies have demonstrated that significantly lower concentrations of GDNF are required to enhance survival, compared to promoting axonal growth and plasticity. In this regard, several regulatory elements are now being tested to control the level of GDNF expression, as well as fine tune duration, that may be necessary for the optimisation of GDNF gene therapy – see review ^10^.

In addition to the support of human pluripotent stem cell derived DA grafts, GDNF gene therapy may provide the supplementary advantage of protecting residual host DA neurons within the VM that are undergoing persistent degeneration. Similar to previous reports ^18, 19^, including larger non-human primates ^21^, and evidenced here, viral delivery of GDNF within the target striatal/caudate putamen tissue also resulted in transport and transduction of cells within the midbrain, as well as activation of downstream MAPK (pERK) and PI3K-AKT (pS6) signalling. Although mature DA neurons have been suggested to be less responsive to GDNF than immature DA progenitors/neurons, studies have provided evidence of improved survival in the adult brain, resulting in a number of clinical trials using GDNF (infusion and viral delivery), as well as the closely associated protein Neurturin, targeted at slowing disease progression ^10, 57^.

The combined approach of cell and gene therapy in neural repair is becoming increasingly recognised for clinical application. Studies have already demonstrated delayed disease progression using human neural progenitors engineered to over-express GDNF in rodent and non-human primate models of PD ^58^, with these approaches also confirmed to be safe in a Phase 1/2a clinical trial targeting protection of motor neurons in ALS (NCT02943850, 2017). Here we demonstrate the benefit of enriching for correctly specified ventral midbrain progenitors, enabling enhanced predictability of graft outcomes, and importantly their responsiveness to targeted GDNF gene therapy. Irrespective of whether reporter lines or alternative cell sorting strategies (such as selection based upon cell surface proteins ^28, 59^) are adopted in the future for clinical application, the impetus remains for multifaceted approaches inclusive of enriching for VM progenitors as well as the implementation of inducible GDNF transgene delivery systems.

In summary, here we provide a conclusive body of evidence for the many benefits of GDNF gene therapy to improve the functionality of human pluripotent stem cell-derived DA neural transplants in rodent models of PD. Dependent of timing of gene delivery, we report that GDNF increases graft survival, plasticity/sprouting of DA fibers, DA innervation of host target nuclei, increased activation of striatal neurons and elevated DA metabolism (associated with changes in regulatory synthesis gene expression). These findings suggest that a dual therapeutic approach, involving cell transplantation and targeted gene therapy, to address the shortcoming of poor survival and plasticity of human DA neurons, may have significant implications for the translation of pluripotent stem cell-based therapies into the clinic for the treatment of Parkinson’s disease.

## Online Methods

### Human ESC Differentiation

The human embryonic stem cell (hESC) reporter lines, H9 PITX3-eGFP and H9 LMX1A-eGFP, were expanded and differentiated as previously described in detail ^2^, and illustrated in Figure 1A. In brief, neural induction was achieved by dual SMAD inhibition using LDN193189 (200 nM, 0–11 days in vitro, DIV; Stemgent) and SB431542 (10 mM, 0–5DIV; R&D Systems). Ventral midbrain patterning was simultaneously achieved by supplementation of the media with sonic hedgehog (200 ng/mL, 1-7DIV; R&D Systems), purmorphamine (2 mM, 1–7DIV; Stemgent) and CHIR99021 (3 mM, 3–13DIV; Miltenyi Biotech). From 11DIV, cultures were matured in the presence of brain-derived neurotrophic factor (BDNF, 20ng/ml, R&D Systems), glial cell line-derived neurotrophic factor (GDNF, 20ng/ml, R&D Systems), recombinant human transforming growth factor type β3 (TGFβ3, 1ng/ml, Peprotech), ascorbic acid (200nM, Sigma-Aldrich), dibutyryl cAMP (0.05mM, Tocris) and DAPT (10μM, Sigma-Aldrich). At 20 DIV cultures enriched with ventral midbrain progenitors were dissociated using Accutase (Life Technologies) in preparation for transplantation.

### Surgical procedures

All animal procedures adhered to the Australian National Health and Medical Research Council’s (NHMRC) Code of Practice for the Use of Animals in Research and were approved by the Florey Institute for Neuroscience and Mental Health animal ethics committee. Animals were group housed on a 12:12-hour light/dark cycle with *ad libitum* access to food and water. Surgeries were performed on 78 athymic (CBH^rnu^) nude rats and 26 athymic (Foxn1^nu^) nude mice under 2-5% isofluorane anaesthesia. Animals received unilateral 6-OHDA lesions (rats: 3.5µl, 3.2μg/μl; mice: 1.5µl, 1.6 µg/µl) of the medial forebrain bundle (MFB; rat) or substantia nigra (mouse), as previously described ^5, 19^.

A subset of animals received an injection of the adeno-associated viral vector carrying GDNF under the chicken ß-actin promoter (AAV-GDNF, 0.25μl; 7.2e^12^ gc/ml) or a control virus, expressing mCherry fluorescent protein (AAV-mCherry, 0.25μl; 4.1e^12^ gc/ml) into the dorsolateral striatum (Rats: 0.5mm rostral, 3.5mm lateral to Bregma and 3.6mm below the surface of the brain; Mice: 1mm rostral, 2mm lateral to Bregma and 2.9mm below the surface of the brain). Human PSC-derived DA progenitors were transplanted into the denervated striatum (1μl; 100,000 cells/μl) (Rats: 0.5mm anterior, 2.5mm lateral to Bregma and 4.0mm below the dural surface; Mice: 0.5mm anterior, 2.5mm lateral, 4.0mm ventral), as previously described ^5, 19^.

#### Study I - Transplantation of progenitors into a GDNF-overexpressing environment

AAV-GDNF (or AAV-mCherry as control) was injected into rats 5 weeks after 6-OHDA lesioning. After a further 3 weeks, these animals were implanted with VM progenitors differentiated from the H9 PITX3-eGFP human ESC line (100,000 cells in 1ul). *Study II – Delayed exposure of the transplant to GDNF*: Eight weeks following 6-OHDA lesioning, rats and mice received implants of 100,000 FACS isolated GFP-expressing VM progenitors, obtained from LMX1A-GFP differentiated hESC cultures. Cell sorting was performed using previously described methods (100μm nozzle at 22PSI, MoFlo XDP (Beckman Coulter)) ^59^. Three weeks later AAV-GDNF was injected into the host striatum. *Study III – Acute assessment of grafts’ responsiveness to GDNF:* 6OHDA lesioned mice received grafts of 100,000 VM progenitors differentiated from the H9 PITX3-eGFP human ESC line, to enable specific tracking of graft-derived (GFP+) DA cells. Three weeks later AAV-GDNF was injected into the host striatum. A subset of animals confirmed expression of the GDNF transgene (by GDNF histochemistry) after 10 days, but not earlier, and hence control and AAV-GDNF animals were killed at day 14, enabling assessment of grafts after acute(~4day) expose to GDNF.

### Behavioural Analysis

Four weeks after lesioning, unilateral DA function was assessed using the amphetamine-induced rotation and cylinder tests, as previously described ^5, 60^, with re-testing performed at various intervals following transplantation. In brief, *Amphetamine rotations*: net rotations over 60 minutes were analysed 10 minutes after intraperitoneal injection of D-amphetamine sulfate (5mg/kg; Tocris Bioscience). *Cylinder test*: animals were placed within a clear glass cylinder and the first 20 forepaw touches were recorded over three consecutive days. Data is presented as left paw ratio: (Left/(Left + Right))*100. Upon completion of initial testing 4 weeks post-lesioning, animals displaying a functional deficit (>300 rotations in 60 min) were ranked in order of the percentage rotational asymmetry and evenly distributed across the four treatment groups (Lesion, Lesion+GDNF, Lesion+Graft, Lesion+Graft+GDNF). The behavioural testing and allocation to (4) treatment groups was performed on 2 cohorts of rats, for Study I and Study II, described above.

### Immunohistochemistry

Animals received intraperitoneal D-amphetamine sulfate (5mg/kg) one hours prior to receiving an overdose of sodium pentobarbitone (100mg/kg) followed by transcardial perfusion with Tryode solution followed by 4% paraformaldehyde. Brains were coronally or horizontally sectioned (40μm; 12 series) on a freezing microtome (Leica) and immunohistochemistry performed as previously described ^60^. Primary antibodies and dilutions are shown in table 1. The bound antibody complex was detected either indirectly through enzymatic conjugation of the diaminobenzadine chromagen or using directly conjugated fluorescent secondary antibodies (Jackson Immunoresearch).

**Table 1.** List of primary antibodies

| **Antibody** | **Species** | **Dilution** | **Source** | **Catalogue #** |
| --- | --- | --- | --- | --- |
| anti-adenomatous polyposis coli,CC1 (APC) | Mouse | 1:100 | Abcam | AB16794 |
| BARHL1 | Rabbit | 1:300 | Novus Biologics | NBP1-86513 |
| Calbindin–C28K | Mouse | 1:1000 | Swant | CB300 |
| Cholecystokinin (CCK) | Rabbit | 1:5000 | P. Frey <sup>63</sup> | N/A |
| cFOS | Goat | 1:1000 | Santa Cruz | sc-52 |
| CTIP2 | Rat | 1:500 | Abcam | AB18465 |
| DAPI | - | 1:5000 | Sigma Aldrich | D8417 |
| DARPP-32 | Rabbit | 1:500 | Millipore | AB10518 |
| FOXA2 | Goat | 1:200 | Santa Cruz | sc-6554 |
| GDNF | Goat | 1:1000 | R&D Systems | AB-212-NA |
| GFAP | Rabbit | 1:500 | DAKO | Z0334 |
| GFP | Chicken | 1:1000 | Abcam | AB13970 |
| GFP | Rabbit | 1:20,000 | Abcam | AB290 |
| GIRK2 | Rabbit | 1:500 | Alomone Labs | APC-006 |
| Human Nuclei (HNA) | Mouse | 1:300 | Millipore | MAB1281 |
| NeuN | Rabbit | 1:100 | R&D Systems | MAB377 |
| NURR1 | Rabbit | 1:200 | Santa Cruz | Sc-990 |
| OTX2 | Rabbit | 1:3000 | Millipore | AB9566 |
| pERK | Rabbit | 1:300 | Cell Signalling Technology | 4370S |
| PITX2 | Sheep | 1:500 | R&D Systems | AF7388 |
| pS6 | Rabbit | 1:300 | Cell Signalling Technology | 2211S |
| PSA-NCAM | Mouse | 1:200 | Santa Cruz | sc-106 |
| RFP | Rabbit | 1:1000 | Rockland labs. | 600-901-379 |
| SYNAPTOPHYSIN (hSYP) | Mouse | 1:1000 | Enzo Life Sciences | ADI-905-782 |
| TH | Rabbit | 1:500 | Pel-freeze | P40101-0 |
| TH | Sheep | 1:800 | Pel-freeze | P60101-0 |
| 5-HT | Rabbit |  | ImmunoStar | 20080 |

### Microscopy and Quantification

Fluorescence images were captured on a Zeiss Axio ObserverZ.1 upright epifluorescence or Zeiss LSM 780 confocal microscope. Bright and darkfield images were taken on a Leica DM6000 upright microscope. Grafts were delineated by human PSA-NCAM expression and volume extrapolated using Cavalieri’s principle. For GFP, TH and cFOS+ quantification, all positive cells were counted from brightfield images and corrected for series number. For GIRK2 and Calbindin quantification, dopamine neurons were first identified using TH immunoreactivity from confocal images. For fiber density assessment, 10 z-stack-sections (1μm per section) were obtained and compressed. TH+ fibers were isolated on colour inverted images using the ‘colour range’ tool on Photoshop (Adobe). Data is expressed as percentage of immunoreactive pixels. All areas were captured in triplicate with conserved settings. Sampling for fiber density and cFOS labelled neurons was performed in the medial (AP: 0.0, ML: −1.8 and DV: −5.1), dorsolateral (AP: 0.0, ML: −3.8 and DV: −3.9) and ventrolateral (AP: 0.0, ML: −4.0 and DV: −6.9) striatum of rats, (from sections 1 mm anterior to 1.5mm posterior to Bregma).

### Gene Expression Analysis

For transcriptional profiling, the ipsilateral striatal hemisphere containing the LMX1A-GFP transplant was dissected from mice 6 months after transplantation. Total RNA was extracted and RNAseq (Illumina) performed. Alignment against the human genome was conducted to select only human specific sequences for analysis using our previously described methods ^33^. GO enrichment analysis (biological process) was performed using the DAVID gene ontology browser, protein-protein interactions were probed using STRING (http://string-db.org/)^41^. A number of genes showing significant change in expression (between Graft and Graft + GDNF) were validated using quantitative real time PCR (qPCR), according to previously described methods ^61^. To selectively assess these gene expression changes in the graft and not host tissue, quantitative PCR (qPCR) primers were designed against regions containing nucleotide base-pair differences between the mouse and human gene. Primers were validated for their selectivity on control mouse (no expression) and human (gene expression) tissue, prior to testing on grafted samples. See table 2 for a list of primers.

**Table 2.**
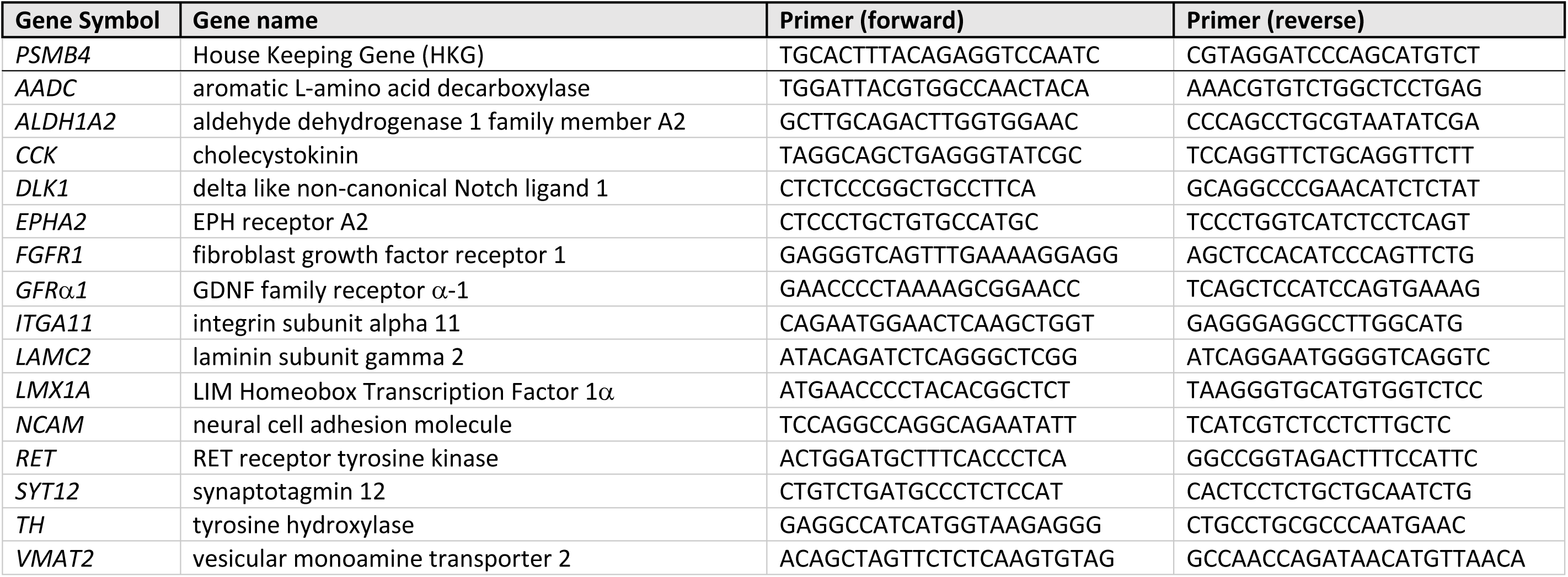
List of primers used for QPCR

Real time PCR, to validate GDNF receptor expression, was performed on undifferentiated PSCs (0 DIV) and differentiated VM progenitors (11, 20 and 25 DIV). The amplicon sizes for GFRa1 and RET were 133 and 122 bp, respectively. In brief, 10ul of the amplified product was loaded per lane and electrophoretically separated using a 3.5% agarose gel at 80V for 55 min.

### HPLC

Using reverse phase liquid chromatography with electrochemical detection, dopamine and its metabolites, 3,4-Dihydroxyphenylacetic acid (DOPAC) and homovanillic acid (HVA), levels were measure from mouse striatal tissue 6 months after the transplantation of LMX1A-GFP VM progenitors, in the absence and presence of GDNF, as previously described ^62^. Data was expressed as pmol/ml of DA, DOPAC or HVA and dopamine turnover determined by the ratio of DA to HVA.

### Statistical Analysis

All data are presented as mean ± SEM. Statistical tests employed (inclusive of one-and two-way ANOVA and student t-tests) and number of independent culture replicates or animals/group are stated in figure legends. Alpha levels of p<0.05 were considered significant with all statistical analysis performed using GraphPad Prism. *p < 0.05, **p < 0.01, ***p < 0.001 and ****p < 0.0001.

### Data availability

Sequencing data from comparative assessment of human PSC-derived VM progenitor grafts in the absence and presence of GDNF will be made available at the time of acceptance of manuscript.

## Acknowledgements

The authors thank Mong Tien and Davor Stanic for expert technical assistance, Björn Anzelius for production of the AAV vectors and acknowledge the support of the flow cytometry facility at the Melbourne Brain Centre and Bioinformatics platform at the Murdoch Children’s Research Institute, Melbourne, Australia. CP was supported by a Senior Medical Research Fellowship provided by the Viertel Charitable Foundation, Australia and subsequent National Health and Medical Research Council Australia (NHMRC) Senior Research Fellowship. CG is supported by an Australian Postgraduate Award. This work was supported by an NHMRC Australia project grant APP1102704 (to CP), Stem Cells Australia (to CP) and Swedish Research Council grant no. 2012-2586 (to DK). The Florey Institute of Neuroscience and Mental Health acknowledges the strong support from the Victorian Government and in particular the funding from the Operational Infrastructure Support Grant.

## Author Contributions

CLP and CWG conceived the experiments and wrote the manuscript; CWG, IDL, JCN, VP, CPH, NM, JAK, CRB, CME, LHT, CLP performed the experiments; CWP, DK, LHT, CLP provided reagents and expertise; CLP, DK provided the funding.

## Competing interests

The authors declare no competing financial interests

## REFERENCES

1. Barker, R.A., Barrett, J., Mason, S.L. & Bjorklund, A. Fetal dopaminergic transplantation trials and the future of neural grafting in Parkinson’s disease. Lancet neurology 12, 84–91 (2013).

2. Niclis, J.C. et al. Efficiently Specified Ventral Midbrain Dopamine Neurons from Human Pluripotent Stem Cells Under Xeno-Free Conditions Restore Motor Deficits in Parkinsonian Rodents. Stem cells translational medicine 6, 937–948 (2017).

3. Kriks, S. et al. Dopamine neurons derived from human ES cells efficiently engraft in animal models of Parkinson’s disease. Nature 480, 547–551 (2011).

4. Kirkeby, A. et al. Generation of regionally specified neural progenitors and functional neurons from human embryonic stem cells under defined conditions. Cell reports 1, 703–714 (2012).

5. Niclis, J.C. et al. A PITX3-EGFP Reporter Line Reveals Connectivity of Dopamine and Non-dopamine Neuronal Subtypes in Grafts Generated from Human Embryonic Stem Cells. Stem Cell Reports 9, 868–882 (2017).

6. Grealish, S. et al. Human ESC-derived dopamine neurons show similar preclinical efficacy and potency to fetal neurons when grafted in a rat model of Parkinson’s disease. Cell stem cell 15, 653–665 (2014).

7. Castilho, R.F., Hansson, O. & Brundin, P. Improving the survival of grafted embryonic dopamine neurons in rodent models of Parkinson’s disease. Progress in brain research 127, 203–231 (2000).

8. Olanow, C.W., Kordower, J.H. & Freeman, T.B. Fetal nigral transplantation as a therapy for Parkinson’s disease. Trends in neurosciences 19, 102–109 (1996).

9. Lin, L.F., Doherty, D.H., Lile, J.D., Bektesh, S. & Collins, F. GDNF: a glial cell line-derived neurotrophic factor for midbrain dopaminergic neurons. Science 260, 1130–1132 (1993).

10. Kirik, D., Cederfjall, E., Halliday, G. & Petersen, A. Gene therapy for Parkinson’s disease: Disease modification by GDNF family of ligands. Neurobiology of disease 97, 179–188 (2017).

11. Bjorklund, A., Rosenblad, C., Winkler, C. & Kirik, D. Studies on neuroprotective and regenerative effects of GDNF in a partial lesion model of Parkinson’s disease. Neurobiology of disease 4, 186–200 (1997).

12. Thompson, L. & Bjorklund, A. Survival, differentiation, and connectivity of ventral mesencephalic dopamine neurons following transplantation. Progress in brain research 200, 61–95 (2012).

13. Rosenblad, C., Martinez-Serrano, A. & Bjorklund, A. Glial cell line-derived neurotrophic factor increases survival, growth and function of intrastriatal fetal nigral dopaminergic grafts. Neuroscience 75, 979–985 (1996).

14. Ahn, Y.H. et al. Increased fiber outgrowth from xeno-transplanted human embryonic dopaminergic neurons with co-implants of polymer-encapsulated genetically modified cells releasing glial cell line-derived neurotrophic factor. Brain Res Bull 66, 135–142 (2005).

15. Sinclair, S.R. et al. GDNF enhances dopaminergic cell survival and fibre outgrowth in embryonic nigral grafts. Neuroreport 7, 2547–2552 (1996).

16. Yurek, D.M. Glial cell line-derived neurotrophic factor improves survival of dopaminergic neurons in transplants of fetal ventral mesencephalic tissue. Experimental neurology 153, 195–202 (1998).

17. Johansson, M., Friedemann, M., Hoffer, B. & Stromberg, I. Effects of glial cell line-derived neurotrophic factor on developing and mature ventral mesencephalic grafts in oculo. Experimental neurology 134, 25–34 (1995).

18. Thompson, L.H., Grealish, S., Kirik, D. & Bjorklund, A. Reconstruction of the nigrostriatal dopamine pathway in the adult mouse brain. Eur J Neurosci 30, 625–638 (2009).

19. Kauhausen, J., Thompson, L.H. & Parish, C.L. Cell intrinsic and extrinsic factors contribute to enhance neural circuit reconstruction following transplantation in Parkinsonian mice. The Journal of physiology 591, 77–91 (2013).

20. Redmond, D.E., Jr. et al. Embryonic substantia nigra grafts in the mesencephalon send neurites to the host striatum in non-human primate after overexpression of GDNF. The Journal of comparative neurology 515, 31–40 (2009).

21. Elsworth, J.D. et al. AAV2-mediated gene transfer of GDNF to the striatum of MPTP monkeys enhances the survival and outgrowth of co-implanted fetal dopamine neurons. Experimental neurology 211, 252–258 (2008).

22. Wakeman, D.R. et al. Human neural stem cells survive long term in the midbrain of dopamine-depleted monkeys after GDNF overexpression and project neurites toward an appropriate target. Stem cells translational medicine 3, 692–701 (2014).

23. Winkler, C., Georgievska, B., Carlsson, T., Lacar, B. & Kirik, D. Continuous exposure to glial cell line-derived neurotrophic factor to mature dopaminergic transplants impairs the graft’s ability to improve spontaneous motor behavior in parkinsonian rats. Neuroscience 141, 521–531 (2006).

24. Georgievska, B., Carlsson, T., Lacar, B., Winkler, C. & Kirik, D. Dissociation between short-term increased graft survival and long-term functional improvements in Parkinsonian rats overexpressing glial cell line-derived neurotrophic factor. Eur J Neurosci 20, 3121–3130 (2004).

25. Piccini, P. et al. Delayed recovery of movement-related cortical function in Parkinson’s disease after striatal dopaminergic grafts. Annals of neurology 48, 689–695 (2000).

26. Kirkeby, A. et al. Predictive Markers Guide Differentiation to Improve Graft Outcome in Clinical Translation of hESC-Based Therapy for Parkinson’s Disease. Cell stem cell 20, 135–148 (2017).

27. Doi, D. et al. Isolation of human induced pluripotent stem cell-derived dopaminergic progenitors by cell sorting for successful transplantation. Stem Cell Reports 2, 337–350 (2014).

28. Samata, B. et al. Purification of functional human ES and iPSC-derived midbrain dopaminergic progenitors using LRTM1. Nature communications 7, 13097 (2016).

29. Hagell, P. et al. Dyskinesias following neural transplantation in Parkinson’s disease. Nat Neurosci 5, 627–628 (2002).

30. Carlsson, T., Carta, M., Winkler, C., Bjorklund, A. & Kirik, D. Serotonin neuron transplants exacerbate L-DOPA-induced dyskinesias in a rat model of Parkinson’s disease. J Neurosci 27, 8011–8022 (2007).

31. de Luzy, I.R. et al. Isolation of LMX1a ventral midbrain progenitors improves the safety and predictability of human pluripotent stem cell-derived neural transplants in Parkinsonian Disease J Neurosci (2019).

32. Cenci, M.A., Kalen, P., Mandel, R.J., Wictorin, K. & Bjorklund, A. Dopaminergic transplants normalize amphetamine-and apomorphine-induced Fos expression in the 6-hydroxydopamine-lesioned striatum. Neuroscience 46, 943–957 (1992).

33. Bye, C.R. et al. Transcriptional profiling of xenogeneic transplants: examining human pluripotent stem cell-derived grafts in the rodent brain. Stem Cell Reports (2019).

34. Jacobs, F.M. et al. Identification of Dlk1, Ptpru and Klhl1 as novel Nurr1 target genes in meso-diencephalic dopamine neurons. Development 136, 2363–2373 (2009).

35. Surmacz, B. et al. DLK1 promotes neurogenesis of human and mouse pluripotent stem cell-derived neural progenitors via modulating Notch and BMP signalling. Stem cell reviews 8, 459–471 (2012).

36. Christophersen, N.S. et al. Midbrain expression of Delta-like 1 homologue is regulated by GDNF and is associated with dopaminergic differentiation. Exp Neurol 204, 791–801 (2007).

37. Kikuchi, T. et al. Human iPS cell-derived dopaminergic neurons function in a primate Parkinson’s disease model. Nature 548, 592–596 (2017).

38. Wang, Y., Perng, S.L., Lin, J.C. & Tsao, W.L. Cholecystokinin facilitates methamphetamine-induced dopamine overflow in rat striatum and fetal ventral mesencephalic grafts. Experimental neurology 130, 279–287 (1994).

39. Agersnap, M., Zhang, M.D., Harkany, T., Hokfelt, T. & Rehfeld, J.F. Nonsulfated cholecystokinins in cerebral neurons. Neuropeptides 60, 37–44 (2016).

40. Hokfelt, T. et al. Evidence for coexistence of dopamine and CCK in meso-limbic neurones. Nature 285, 476–478 (1980).

41. Szklarczyk, D. et al. The STRING database in 2017: quality-controlled protein-protein association networks, made broadly accessible. Nucleic Acids Res 45, D362–D368 (2017).

42. Kramer, E.R. & Liss, B. GDNF-Ret signaling in midbrain dopaminergic neurons and its implication for Parkinson disease. FEBS Lett 589, 3760–3772 (2015).

43. Steinbeck, J.A. et al. Optogenetics enables functional analysis of human embryonic stem cell-derived grafts in a Parkinson’s disease model. Nature biotechnology 33, 204–209 (2015).

44. Chang, J.W., Wachtel, S.R., Young, D. & Kang, U.J. Biochemical and anatomical characterization of forepaw adjusting steps in rat models of Parkinson’s disease: studies on medial forebrain bundle and striatal lesions. Neuroscience 88, 617–628 (1999).

45. Mandel, R.J., Brundin, P. & Bjorklund, A. The Importance of Graft Placement and Task Complexity for Transplant-Induced Recovery of Simple and Complex Sensorimotor Deficits in Dopamine Denervated Rats. Eur J Neurosci 2, 888–894 (1990).

46. Piccini, P. et al. Factors affecting the clinical outcome after neural transplantation in Parkinson’s disease. Brain 128, 2977–2986 (2005).

47. Carlsson, T. et al. Graft placement and uneven pattern of reinnervation in the striatum is important for development of graft-induced dyskinesia. Neurobiology of disease 21, 657–668 (2006).

48. Grealish, S. et al. The A9 dopamine neuron component in grafts of ventral mesencephalon is an important determinant for recovery of motor function in a rat model of Parkinson’s disease. Brain 133, 482–495 (2010).

49. Borgal, L., Hong, M., Sadi, D. & Mendez, I. Differential effects of glial cell line-derived neurotrophic factor on A9 and A10 dopamine neuron survival in vitro. Neuroscience 147, 712–719 (2007).

50. Kholodilov, N. et al. Regulation of the development of mesencephalic dopaminergic systems by the selective expression of glial cell line-derived neurotrophic factor in their targets. J Neurosci 24, 3136–3146 (2004).

51. Nosheny, R.L., Bachis, A., Aden, S.A., De Bernardi, M.A. & Mocchetti, I. Intrastriatal administration of human immunodeficiency virus-1 glycoprotein 120 reduces glial cell-line derived neurotrophic factor levels and causes apoptosis in the substantia nigra. J Neurobiol 66, 1311–1321 (2006).

52. Reyes, S. et al. GIRK2 expression in dopamine neurons of the substantia nigra and ventral tegmental area. The Journal of comparative neurology 520, 2591–2607 (2012).

53. Xiao, H., Hirata, Y., Isobe, K. & Kiuchi, K. Glial cell line-derived neurotrophic factor up-regulates the expression of tyrosine hydroxylase gene in human neuroblastoma cell lines. Journal of neurochemistry 82, 801–808 (2002).

54. Eslamboli, A. et al. Continuous low-level glial cell line-derived neurotrophic factor delivery using recombinant adeno-associated viral vectors provides neuroprotection and induces behavioral recovery in a primate model of Parkinson’s disease. J Neurosci 25, 769–777 (2005).

55. Kordower, J.H. et al. Neurodegeneration prevented by lentiviral vector delivery of GDNF in primate models of Parkinson’s disease. Science 290, 767–773 (2000).

56. Akhtar, A.A. et al. Inducible Expression of GDNF in Transplanted iPSC-Derived Neural Progenitor Cells. Stem Cell Reports 10, 1696–1704 (2018).

57. Rangasamy, S.B., Soderstrom, K., Bakay, R.A. & Kordower, J.H. Neurotrophic factor therapy for Parkinson’s disease. Progress in brain research 184, 237–264 (2010).

58. Behrstock, S. et al. Human neural progenitors deliver glial cell line-derived neurotrophic factor to parkinsonian rodents and aged primates. Gene Ther 13, 379–388 (2006).

59. Bye, C.R., Jonsson, M.E., Bjorklund, A., Parish, C.L. & Thompson, L.H. Transcriptome analysis reveals transmembrane targets on transplantable midbrain dopamine progenitors. Proceedings of the National Academy of Sciences of the United States of America 112, E1946–1955 (2015).

60. Somaa, F.A. et al. Peptide-Based Scaffolds Support Human Cortical Progenitor Graft Integration to Reduce Atrophy and Promote Functional Repair in a Model of Stroke. Cell reports 20, 1964–1977 (2017).

61. Somaa, F.A., Bye, C.R., Thompson, L.H. & Parish, C.L. Meningeal cells influence midbrain development and the engraftment of dopamine progenitors in Parkinsonian mice. Experimental neurology 267, 30–41 (2015).

62. Parish, C.L. et al. Wnt5a-treated midbrain neural stem cells improve dopamine cell replacement therapy in parkinsonian mice. The Journal of clinical investigation 118, 149–160 (2008).

63. Frey, P. Cholecystokinin octapeptide levels in rat brain are changed after subchronic neuroleptic treatment. Eur J Pharmacol 95, 87–92 (1983).

