## supplementary data for "VIRAL DELIVERY OF GDNF PROMOTES FUNCTIONAL INTEGRATION OF HUMAN STEM CELL GRAFTS IN PARKINSON’S DISEASE"

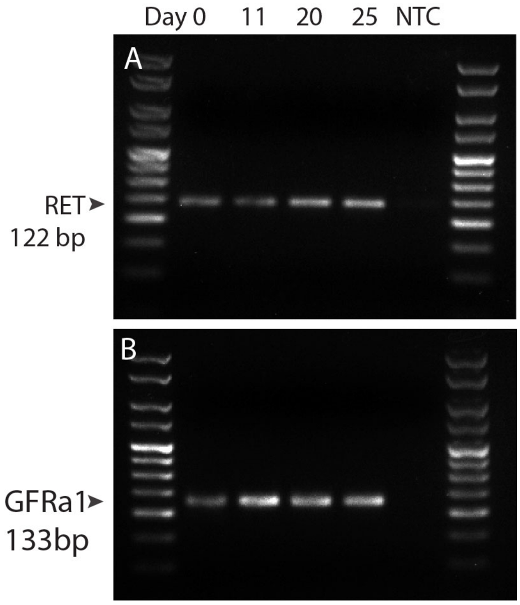


**Supplementary Figure 1: RT-PCR confirmation of GDNF receptor expression on transplanted progenitors.**

(A-B) Representative gels confirming expression of (A) Ret and (B) GFRa1 on the undifferentiated PSCs and VM progenitors (at 11, 20 and 25 days in vitro, DIV). Cell were transplanted at 20-21 DIV.

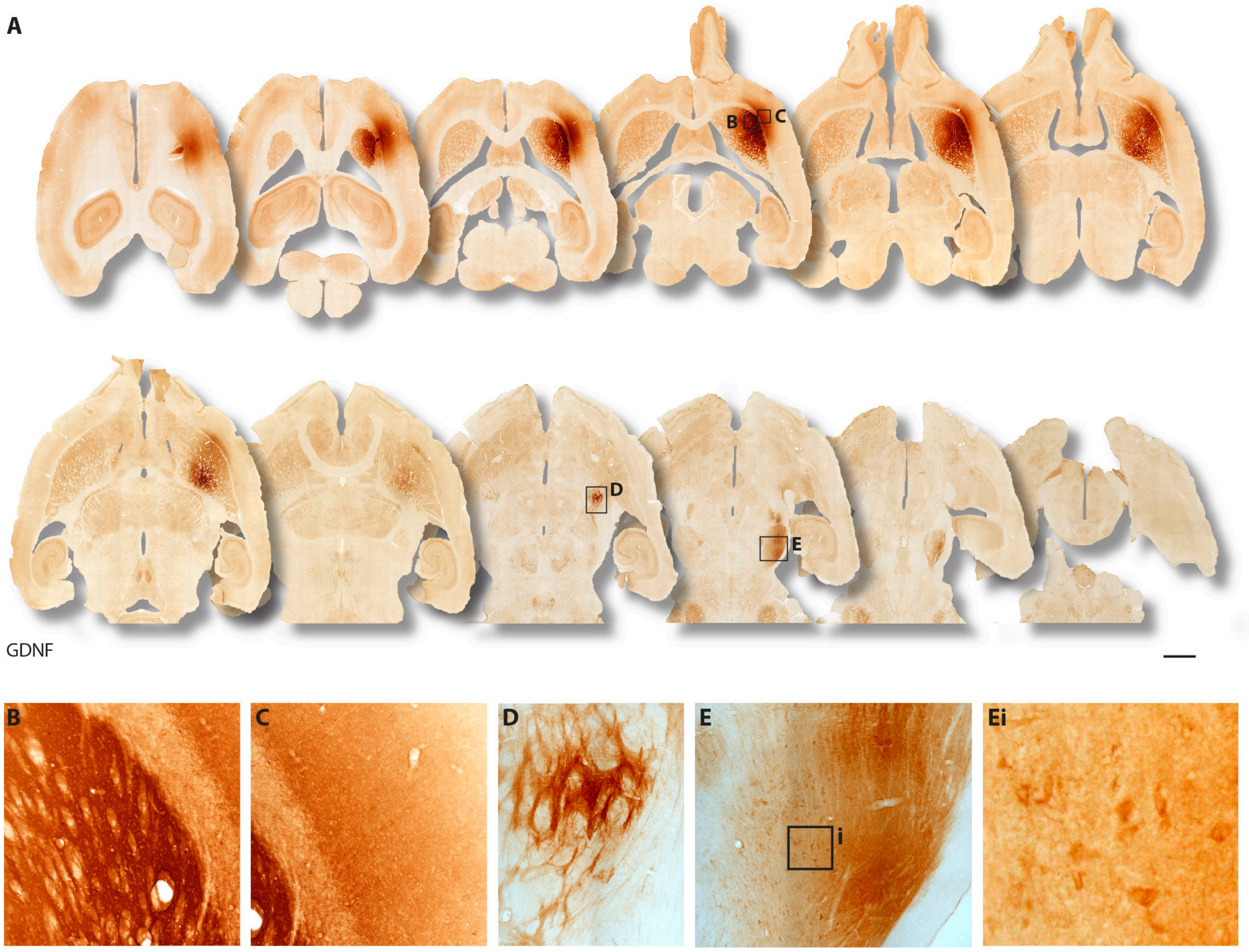


**Supplementary Figure 2: Targeted delivery of AAV-GDNF into the host brain. genes.**
(A) GDNF immunohistochemistry confirmed sustained expression of the protein within the host tissue 6 months after intrastriatal injection. In addition to expression within the host striatum and overlying cortex, GDNF was present within the MFB and VM. High power photomicrographs confirmed viral transduction of cells within the (B) striatum, (C) overlying cortex, (D) MFB and (E) VM cells. Scale bar, 2mm.

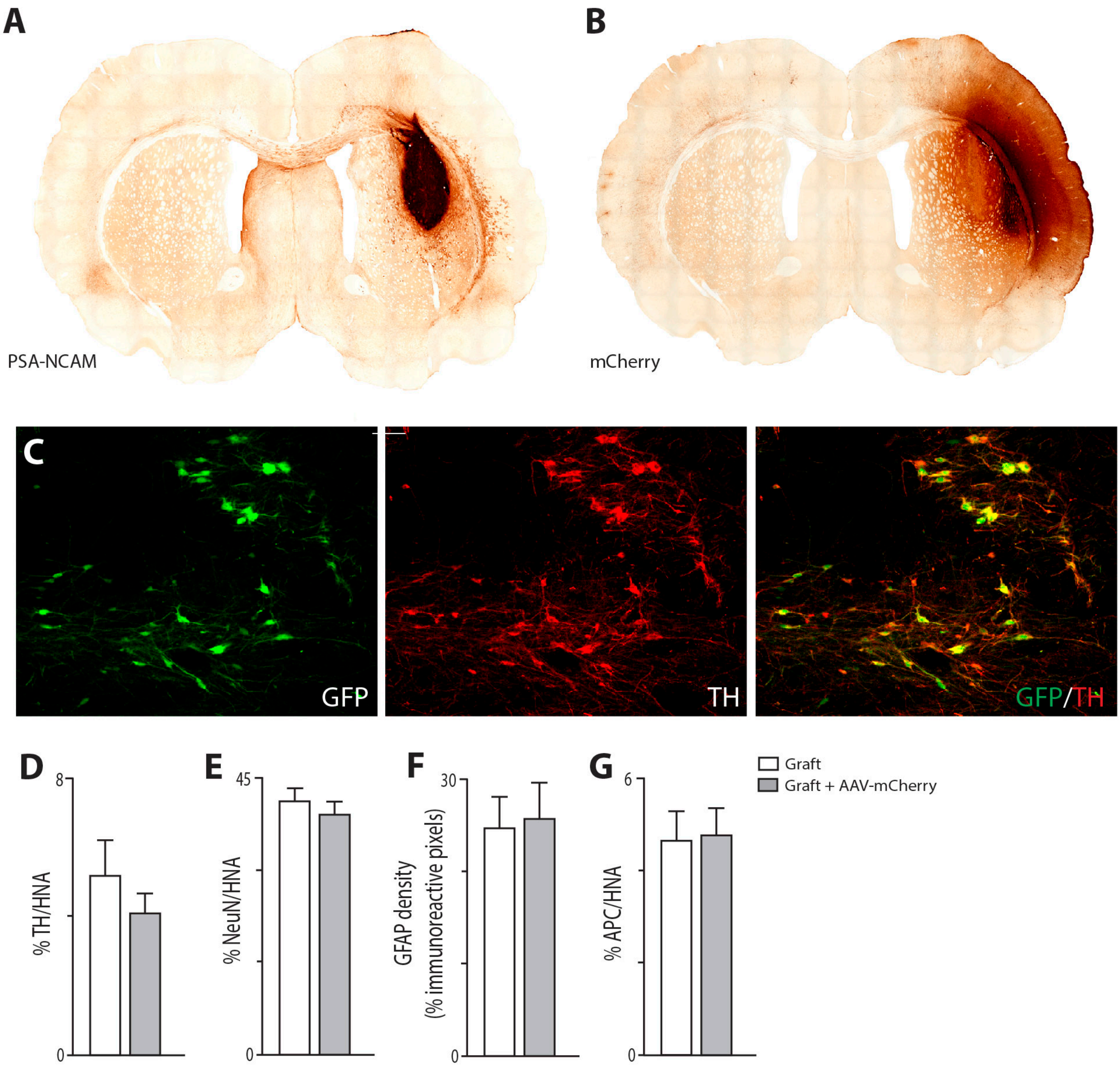


**Supplementary Figure 3: Control AAV-mCherry has no effect on the survival or differentiation of hESC-derived VM progenitor grafts**

(A) Representative image of PSA-NCAM immunoreactivity delineating a hESC-derived PITX3-eGFP graft in the presence of the control AAV-mCherry.

(B) Photomicrograph illustrating the targeted localisation of the mCherry virus to the host striatum and overlying cortex.

(C) AAV5-mCherry had no effect on the ability of transplanted progenitors to differentiate into dopaminergic neurons as assessed by GFP+ (PITX3-GFP) and TH+ co-expression.

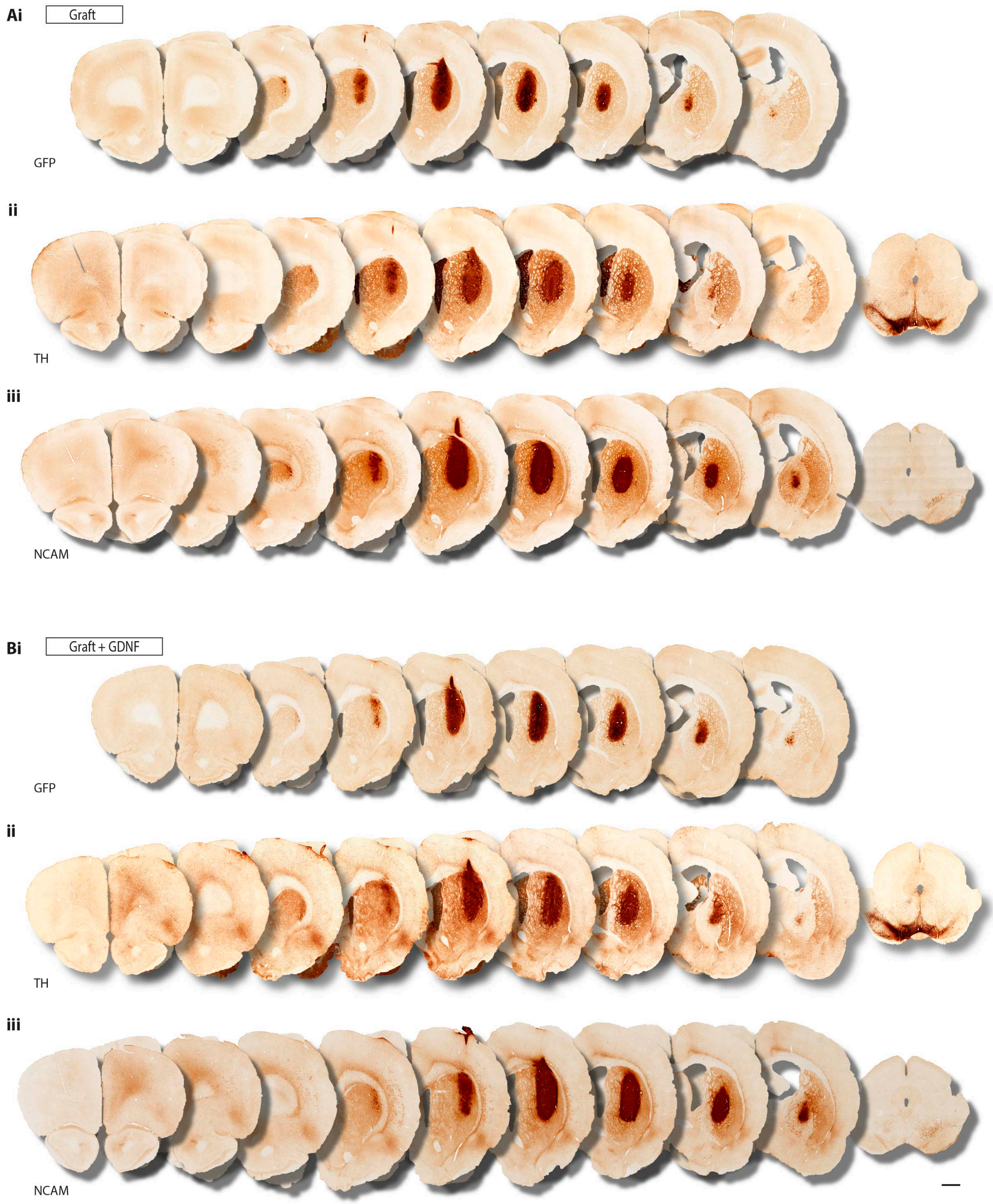


**Supplementary Figure 4: Comparative assessment of LMX1A-eGFP, TH and NCAM immunoreactivity demonstrates that the vast majority of the transplant, in the absence or presence of GDNF, is DA at 6 months.**

(A) Human ESC-derived VM progenitor grafts in the absence and (B) presence of GDNF, illustrating maintained GFP expression of the early progenitor marker LMX1A (Ai, Bi), that largely overlaps with the dopaminergic neuronal marker tyrosine hydroxylase (TH; Aii,Bii), as well as the human specific antibody for the neural cell adhesion molecule NCAM (Aiii, Biii). Images shown in A (Ai, Aii and Aiii) are taken from adjacent serial sections of the same brain. Similarly, Images shown in Bi, Bii and Biii are from adjacent serial sections. Collectively these images highlight that a major proportion of the transplant was dopaminergic. Scale bar, 1mm.

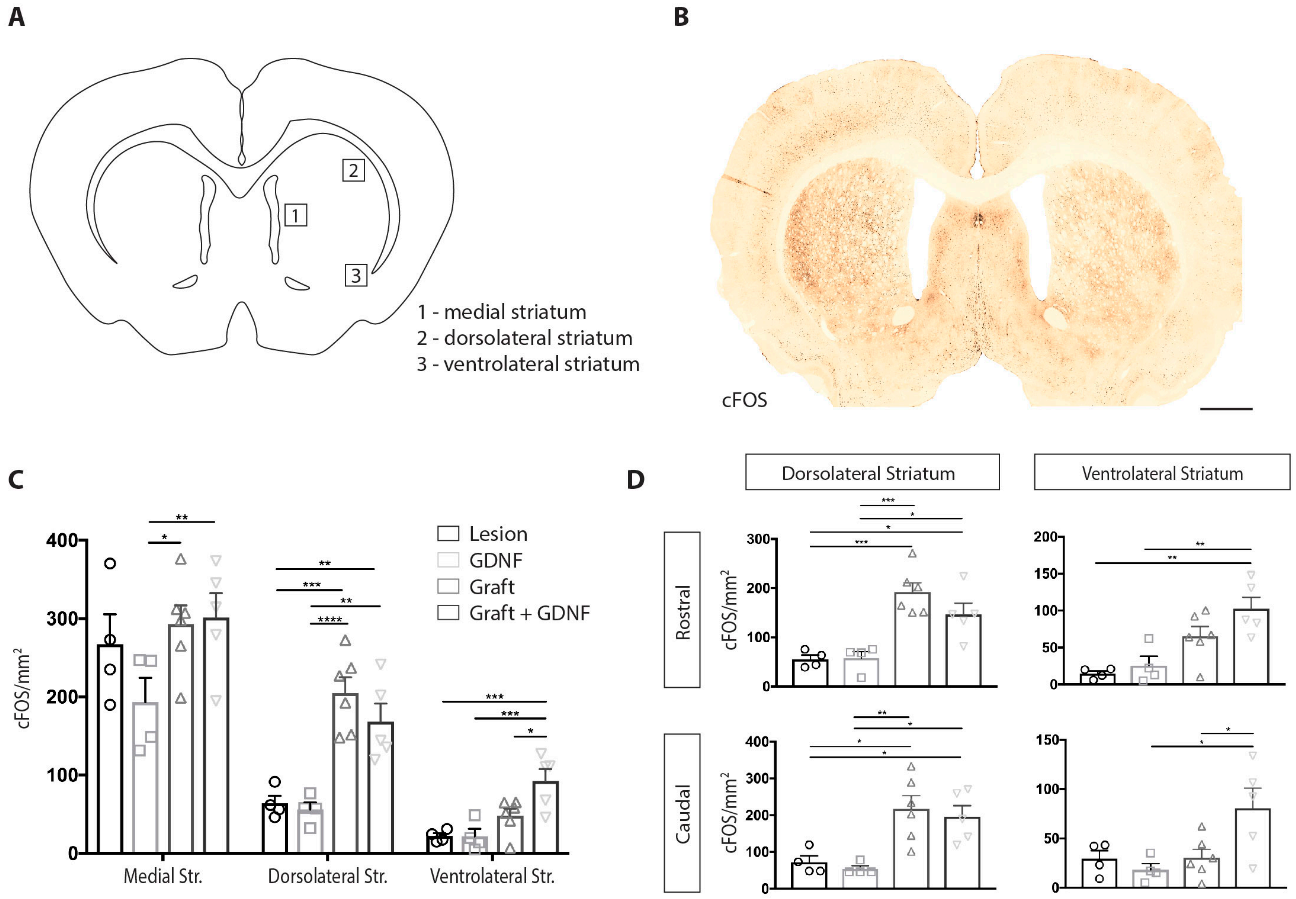


**Supplementary Figure 5: Quantitative analysis of c-FOS activated striatal cells by human PSC-derived VM transplants**

(A) Schematic coronal view of the rat brain illustrating the three sampling sites for quantification of cFOS-immunoreactive cells.

(B) Example photomontage showing diffuse c-FOS staining in dopaminergic target regions of the rat brain following acute amphetamine administration in a 6-OHDA lesioned animal. Scale bar, 1mm.

(C) Quantification of c-FOS density within the medial, dorsolateral and ventrolateral striatum of lesion and grafted animals (± AAV-GDNF). Densities are estimated from 6 serial sections, spanning ~3mm, through the striatum. Two-way ANOVA with Tukey correction for multiple comparisons (n = 4-6).

(D) Quantification of c-FOS+ cell density within the rostral and caudal regions of the dorsolateral and ventrolateral striatum. Densities are estimates taken from 3 sections spanning ~1.5mm through the striatum. One-way ANOVA with Tukey correction for multiple comparisons (n = 4-6).

Abbreviations: Str., Striatum.

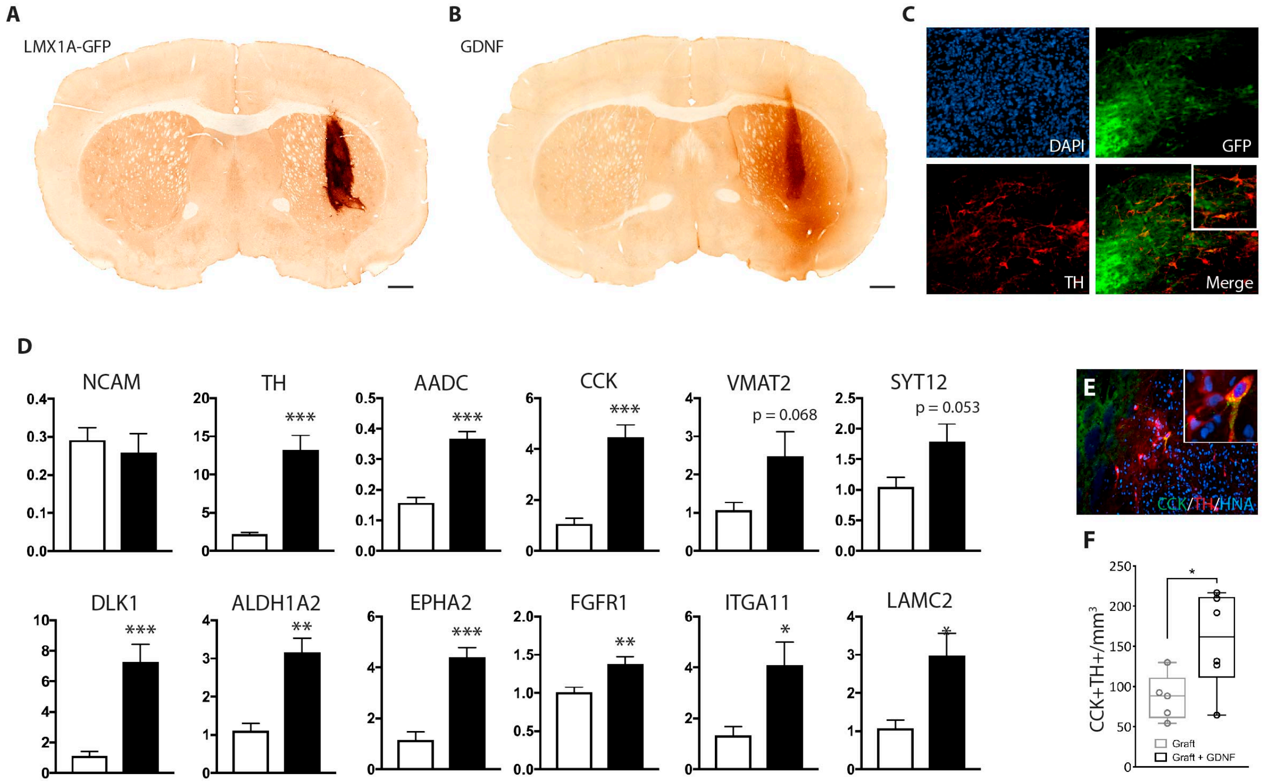


**Supplementary Figure 6: LMX1A-GFP+ human PSC-derived VM progenitor graft in the Parkinsonian mouse brain.**

(A) Representative image of an LMX1A-GFP+ human VM progenitor graft at 6 months in the lesioned nude mouse. LMX1A-GFP grafts in the mouse brain resembled those in the rat brain and were collected for RNAseq and HPLC analysis.

(B) GDNF immunohistochemistry to confirm AAV-GDNF induced overexpression in the mouse brain.

(C) LMX1A-GFP+ grafts matured into TH+ DA neurons in the mouse brain.

(D) Species-specific qPCR of human mRNA expression levels within the graft confirmed the expression of genes identified by RNAseq. In addition to verifying the maintained graft size (unchanged NCAM level) these results supported increases in DA synthesis, plasticity and synaptic protein associated genes as well as extracellular matrix in response to GDNF.

(E) CCK+ neurons were readily identified within the graft and often coexpressed TH.

(F) Quantification of CCK+ and CCK+/TH+ cells within the grafts revealed that both CCK+ and CCK+/TH+ neuronal density was increased in the presence of GDNF.

See table 2 for gene abbreviations.

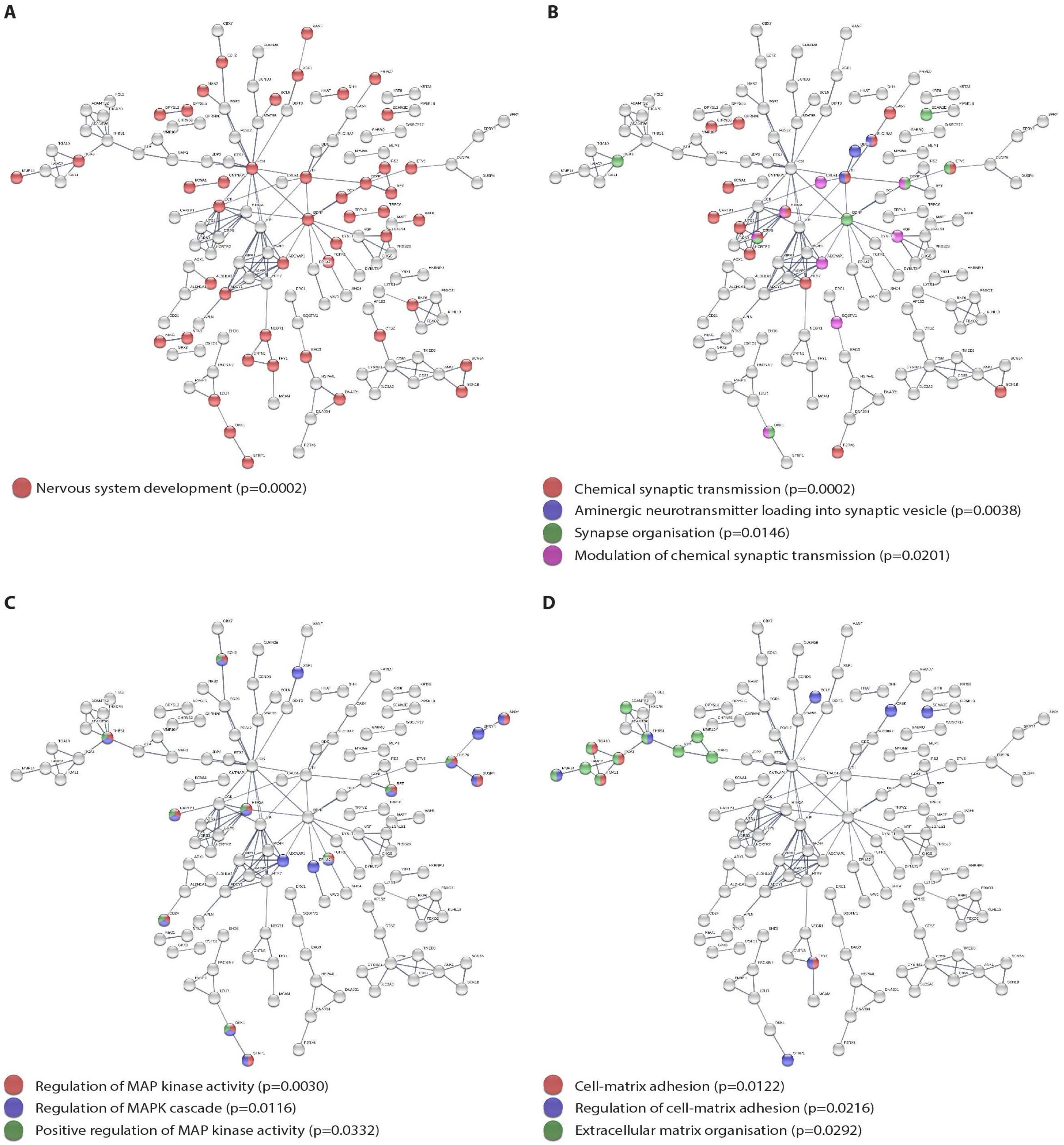


**Supplementary Figure 7: Protein-protein interaction analysis of genes differentially expressed following GDNF treatment.**

(A-D) STRING analysis of differentially expressed genes. Genes were associated with GO terms including nervous system development (A), chemical synaptic transmission (B), MAPK cascades (C) and adhesion (D). Minimum confidence scores of 0.700 were used, line thickness represents confidence score and each GO term is represented by the colour of each gene.
